# STING–STAT3–SOX18 Axis Drives EndMT and Epigenetic Reprogramming in SAVI Lung Fibrosis

**DOI:** 10.64898/2026.03.23.713256

**Authors:** Dan Yang, Guibin Chen, Sachin Gaurav, Adriana A. de Jesus, Atul K. Mehta, Colton McNinch, Adam X. Miranda, Jinzhi Wei, Noemi Kedei, Maria Olga Hernandez, Jizhong Zou, Kaari Linask, Chyi-Chia Lee, Gauthaman Sukumar, Ying Zhang, Sara Alehashemi, Les Folio, Quan Yu, Bin Lin, Bianca Lang, Bjoern Buehring, Gregor Dueckers, Adam Reinhardt, Gregory Schulte, Deborah R. Liptzin, Seza Ozen, Arturo Borzutzky, Madeline Wong, Desiree Tillo, Natalia I. Dmitrieva, Haresh Mani, Steven D. Nathan, Jason C. Kovacic, Clifton Dalgard, Manfred Boehm, Raphaela Goldbach-Mansky

## Abstract

A high prevalence of early-onset interstitial lung disease, including pulmonary fibrosis, in pediatric patients with Stimulator of interferon genes (STING)-Associated Vasculopathy with onset in infancy (SAVI) suggests a critical role for the cGAS-STING pathway in the pathogenesis of pulmonary fibrosis. We identified an endothelial-to-mesenchymal transition (EndMT) signature in lesional lung biopsies from SAVI patients, marked by a loss of endothelial and acquisition of mesenchymal markers. Consistently, induced pluripotent stem cell-derived endothelial cells (iECs) from SAVI patients harboring gain-of-function *STING1* mutations spontaneously undergo EndMT, a process rescued in isogenic-correction. In endothelial cells, STING activation induces IRF3-independent STAT3 phosphorylation, initiating a SLUG-dependent mesenchymal transcriptional program while repressing SOX18 and an epigenetically-regulated endothelial maintenance network. Our studies define a non-canonical cGAS-STING-STAT3 signaling axis that couples a mesenchymal transcriptional program with epigenetic silencing of an endothelial maintenance program, promoting TGFβ-independent STING-mediated EndMT and endothelial dysfunction, and suggesting STING as a therapeutic target for inflammatory pulmonary fibrosis.

## INTRODUCTION

Mendelian autoinflammatory diseases with early-onset organ involvement have revealed critical pathways of innate immune dysregulation that drive chronic inflammation and maladaptive tissue remodeling ^1^. STING-associated vasculopathy with onset in infancy (SAVI), caused by gain-of-function mutations in the nucleic acid sensor *STING1* is a prototypic interferonopathy characterized by severe childhood-onset interstitial lung disease, pulmonary fibrotic lesions leading to respiratory failure that is a major cause of early mortality ^2,3^. Among type I interferonopathies, interstitial lung disease (ILD) is a prominent feature only in SAVI and Coatomer-associated protein subunit alpha (COPA) syndrome, both of which are linked to STING pathway dysregulation. The interstitial lung disease (ILD) in SAVI shares features with systemic autoimmune rheumatic disease–associated ILDs (SARD-ILD), but lacks the fibroblastic foci seen in idiopathic pulmonary fibrosis (IPF), suggesting a unique contribution of STING signaling to the development of lung fibrosis ^4,5^.

Chronic inflammation in the lung can impair tissue regeneration and promote fibrosis by disrupting epithelial and endothelial homeostasis. During tissue repair, multiple cell types, including fibroblasts, epithelial, and endothelial cells, activate a mesenchymal program to support extracellular matrix remodeling ^6,7^. However, persistence of these reparative processes leads to excessive matrix deposition, fibrosis, and ultimately organ failure. The mechanisms that initiate mesenchymal remodeling program in lung fibrosis remain incompletely defined and may vary by disease context.

STING is expressed in multiple pulmonary residential cell lines, including endothelial cells, macrophages, and pneumocytes ^8^ and STING gain-of-function (GOF) mutations in knock-in mice recapitulate SAVI lung pathology independently of type I interferon or IRF3 signaling ^9–11^. Disease in these murine models is not ameliorated by interferon-alpha/beta receptor (IFNAR) blockade or bone marrow transplantation, implicating non-hematopoietic tissue cells in disease initiation ^12^. These findings, together with the poor outcomes of lung transplantation in SAVI patients ^13^, underscore the need to define cell-intrinsic pathways that drive pulmonary fibrosis in SAVI patients.

Histological analyses of lung and skin biopsies from patients with SAVI reveal prominent endothelial injury and vascular remodeling. These findings led us to hypothesize that the cyclic GMP-AMP synthase (cGAS)/STING pathway disrupts endothelial homeostasis and contributes to early pathogenic changes in SAVI lung disease. Endothelial-to-mesenchymal transition (EndMT) is a fundamental process during development ^14,15^ and wound healing ^16,17^ and is increasingly recognized as contributor to tissue fibrosis but remains poorly characterized in inflammatory lung disease ^18–21^. In this study, we identify cGAS/STING activation as a trigger of TGFβ-independent, STAT3/SLUG/SOX18-driven EndMT, and demonstrate that STING signaling represses transcriptional and epigenetic programs essential for endothelial identity. Our findings highlight a mechanistically targetable axis of fibrosis initiation in SAVI and raise the question of a role of STING-mediated pathology in other forms of inflammatory pulmonary fibrosis.

## RESULTS

### Histopathological features of pulmonary fibrosis associated with SAVI reveal microvascular damage and fibrosis

Clinical and demographic data from 13 genetically confirmed SAVI patients revealed that 10 had interstitial lung disease findings on their chest CT with ground glass opacities, cystic changes, reticular opacities (interlobular and intralobular septal thickening), and early and advanced fibrotic changes (including fibrotic reticulations, traction bronchiectasis, volume loss, and honeycombing) (Figure 1A, Figure S1A), which was histologically confirmed in 7 of 10 patients with available lung biopsy specimen (Supplemental Table 1). Six patients harbored the p.N154S *STING1* mutation, three had p.V155M, one carried p.V147L, one p.V147M, one p.H72N, and one patient harbored a biallelic p.R281W variant (Supplemental Table 1). Lung biopsies showed prominent microvascular changes, with increased collagen and elastin deposition, alveolar wall thickening, and perivascular fibrosis (Figure 1B). Inflammatory and fibrotic scores were significantly higher in SAVI patients compared to controls (Figure 1B), and mesenchymal marker SMA/*ACTA2* and a decrease in the endothelial marker CD31/*PECAM* (Figure S1B) showed significant fibrosis with loss of endothelial cells. SAVI patients showed extensive lung remodeling reminiscent of systemic autoimmune rheumatic disease-associated interstitial lung disease (SARD-ILD) including prominent alveolar and vascular changes and interlobular septal thickening, and chronic inflammatory cell infiltration (Figure 1C), but lacked the fibroblastic foci that are a hallmark of idiopathic pulmonary fibrosis (IPF) (Figure S1C). To investigate early changes in alveolar microarchitecture we used multiplex staining by co-detection by indexing (CODEX) ^22^ and compared areas of mild, moderate and severe fibrosis that were selected by lung pathologists blinded to diagnosis (Figure 1D-E, Figure S1D). Areas of mild fibrosis representing early alveolar changes, showed a relative increase in epithelial cells (I) that were E-cadherin positive (II), with predominance of alveolar type 2 cells (AT2) (III) and a decreased endothelial to AT2 cell ratio (IV) suggestive of early AT2 hyperplasia ^23^ (Figure 1E). Areas of moderate and severe fibrosis showed an increase in SMA positive cells (V) with endothelial cells (CD34+) also expressing SMA (CD34+ and SMA+ double positive (VI) Figure 1E). These findings were confirmed by immunofluorescence labeling with the additional endothelial adhesion markers VE-cadherin/*CDH5* and CD31/*PECAM1*, co-stained with SMA/*ACTA2* (Figure 1F, Figure S1E), suggesting an EndMT process in lung tissues from patients with SAVI,

**Fig1:**
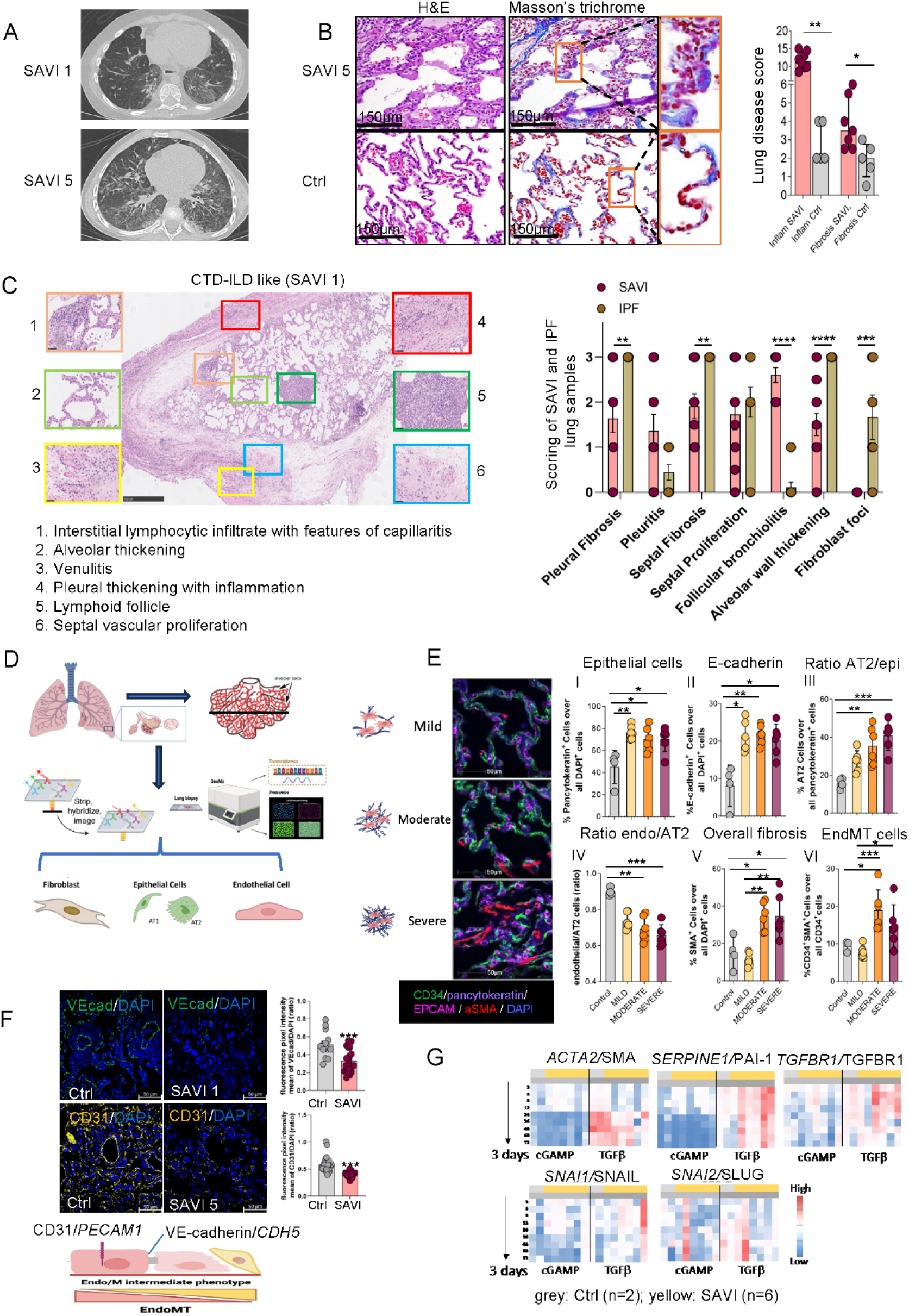
Histopathological features of pulmonary fibrosis associated with SAVI reveal microvascular damage and fibrosis. **A**. Pulmonary CT scans from SAVI patients (SAVI 1 and SAVI 5) showing fibrotic regions. Additional scans are shown in Figure S1A. **B**. Pathological characterization of SAVI lung tissue. Left panels: H&E and Masson-Trichrome staining of alveolar regions (SAVI 5) and controls (Ctrl), highlighting alveolar wall thickening and alveolar capillary fibrosis (orange box and magnified insets). Scale bar: 150 µm. Right panel: Quantification of inflammation and fibrosis scores (n=7 SAVI patients, n=5 controls). Mann-Whitney test; *, 0.01<p<0.05; **, 0.001<p<0.01. (CD31 and SMA staining in Figure S1B). **C**. Pathological features of SAVI-associated interstitial lung disease (11 biopsies from 8 patients) with corresponding higher-magnification insets were compared with idiopathic pulmonary fibrosis (IPF) biopsies (right panel; n = 7). Scored features are listed below; unlike IPF, SAVI lungs lacked fibroblast foci (see also Figure S1C). **D**. CODEX and GeoMx DSP workflow used in this study: paraffin lung sections from controls and SAVI patients were stained with multiplex DNA-conjugated antibodies, imaged, and computationally processed for CODEX. Data were analyzed in regions spanning normal to severe fibrosis (adapted from Nature Protocols 2021 ^22^). FFPE samples from control and SAVI patients were hybridized with probes detecting the whole transcriptome and selected proteins. Regions of interest were selected based on the morphology staining for nuclei, CD45, CD68 and aSMA. Samples were collected from whole ROIs as well as masked regions enriched for aSMA and CD68, and sequenced and analyzed according to vendor’s protocol to detect differences in transcript levels. **E**. Representative CODEX images and quantification of endothelial, epithelial, mesenchymal, and EndMT cells in SAVI lung tissue. Mild, moderate, and severe fibrotic regions in Patient SAVI 8 were identified by Masson’s trichrome staining, and six areas per region were analyzed (Figure S1D). Left panel shows CD34 (green), pancytokeratin (blue-purple), EPCAM (pink-purple), SMA (red), and DAPI (blue) staining (scale bar: 50 µm). SAVI lungs show increased ancytokeratin⁺ epithelial cells (I) and elevated E-cadherin (II). EPCAM⁺ AT2 cell frequency is increased among epithelial cells (III), whereas the CD34⁺ endothelial-to-AT2 ratio is reduced (IV). SMA⁺ mesenchymal cells and CD34⁺/SMA⁺ EndMT cells are significantly increased in moderate and severe fibrosis (V, VI). **F**. Immunofluorescent staining of endothelial markers VE-cadherin (green) and CD31 (yellow) in lung sections from controls (n = 4–5) and SAVI patients (n = 3). Nuclei are labeled with DAPI (blue); scale bar: 50 µm. Mean fluorescence intensity was quantified using ZEN. Both endothelial markers were significantly reduced in SAVI tissue (mean ± SEM; ***p < 0.001, two-tailed unpaired t-test). A schematic of EndMT is shown below; higher-magnification VE-cad/SMA double staining is provided in Figure S1E. **G**. qPCR heatmap of cGAMP- and TGFβ-induced responses in fibroblasts from two controls (grey) and six SAVI patients (yellow). Primary fibroblast cell lines were stimulated with 2’3’-cGAMP or TGFβ for 3–72 hours, and RNA was collected across timepoints to measure mesenchymal gene expression (*ACTA2, SNAI1, SNAI2, SERPINE1, TGFBR1*). Primer and method details are provided in the STAR Methods.

We next assessed the impact of the disease-causing STING mutations on fibroblast activation. Primary dermal fibroblast cell lines from 2 healthy volunteers and 6 SAVI patients were stimulated with cGAMP or TGFβ, the latter known to induce a myofibroblast gene-signature upon activation. Cultured fibroblasts from SAVI patients were not constitutively active but TGFβ stimulation increased myofibroblast markers, including *SERPINE1*/PAI-1 and *ACTA2*/SMA ^24^, similarly in fibroblasts from SAVI patients and healthy controls; stimulation with the STING activator, 2’3’-cGAMP, did not induce myofibroblast markers in SAVI fibroblasts (Figure 1G). These findings suggest that STING mutations do not prime for myofibroblasts differentiation and did not increase TGFβ mediated myofibroblast activation.

Overall, these histological evaluations in lung biopsies from SAVI patients are suggestive that EndMT is a prominent early feature of SAVI-associated pulmonary fibrosis.

### iPSC-derived endothelial cells (iECs) from SAVI patients but not from isogenic and healthy control iECs spontaneously undergo EndMT upon cell passage

To further study EndMT in SAVI, we established an in-vitro disease model described in Figure 2A. Briefly, we generated induced pluripotent stem cells (iPSCs)-derived endothelial cells (iECs) from 4 healthy controls (HC), 4 SAVI patients (3 with *p.N154S* and one with a *p.V147L* mutation) and isogenic control lines (iso-SAVI) parentally from 2 SAVI patients (SAVI 1 and SAVI 2) using CRISPR/Cas9 technology to control for genetic background effects (Figure 2A, Figure S2A). Mutant and wildtype iPSCs similarly differentiated into cuboidal-shaped iECs that express CD31 and CD144 (VE-cadherin), and were morphologically and functionally similar to controls (iECs at passage 1 (P1). (Figure S2B-D). However, in contrast to HC and iso-SAVI iECs, preceding the phenotypic transition around passage 3 (P3) to passage 5 (P5), SAVI iECs progressively lose endothelial surface markers (CD144 (VE-cadherin)/*CDH5* and CD31/*PECAM1)* (Figure 2B, Figure S2C) and develop a spindle-like morphology with impaired angiogenic properties including tube formation (Figure 2C) and decreased EC functional Acetyl-LDL uptake (Figure S2D). Concomitantly, increased protein expression of mesenchymal markers including SMA/*ACTA2* and SM22 (Transgelin)/*TAGLN* in SAVI iECs compared to HC and iso-SAVI iECs at P5 is suggestive of EndMT (Figures 2D-E). STING shRNA knockdown preserved EC surface markers in SAVI iECs, further indicating a STING-dependent signaling pathway (Figure S2E). Moreover, a similar EndMT phenotypic change was observed in primary human lung microvascular endothelial cells (HLMECs) treated with the STING agonist, 2’3’ cyclic guanosine monophosphate (2′3′-cGAMP), which was accelerated when combined with recombinant interferon beta (IFNβ) by day 3 to 5 (Figure S2F-G). In summary, two distinct EC models demonstrated that activation of STING can induces EndMT.

**Fig2:**
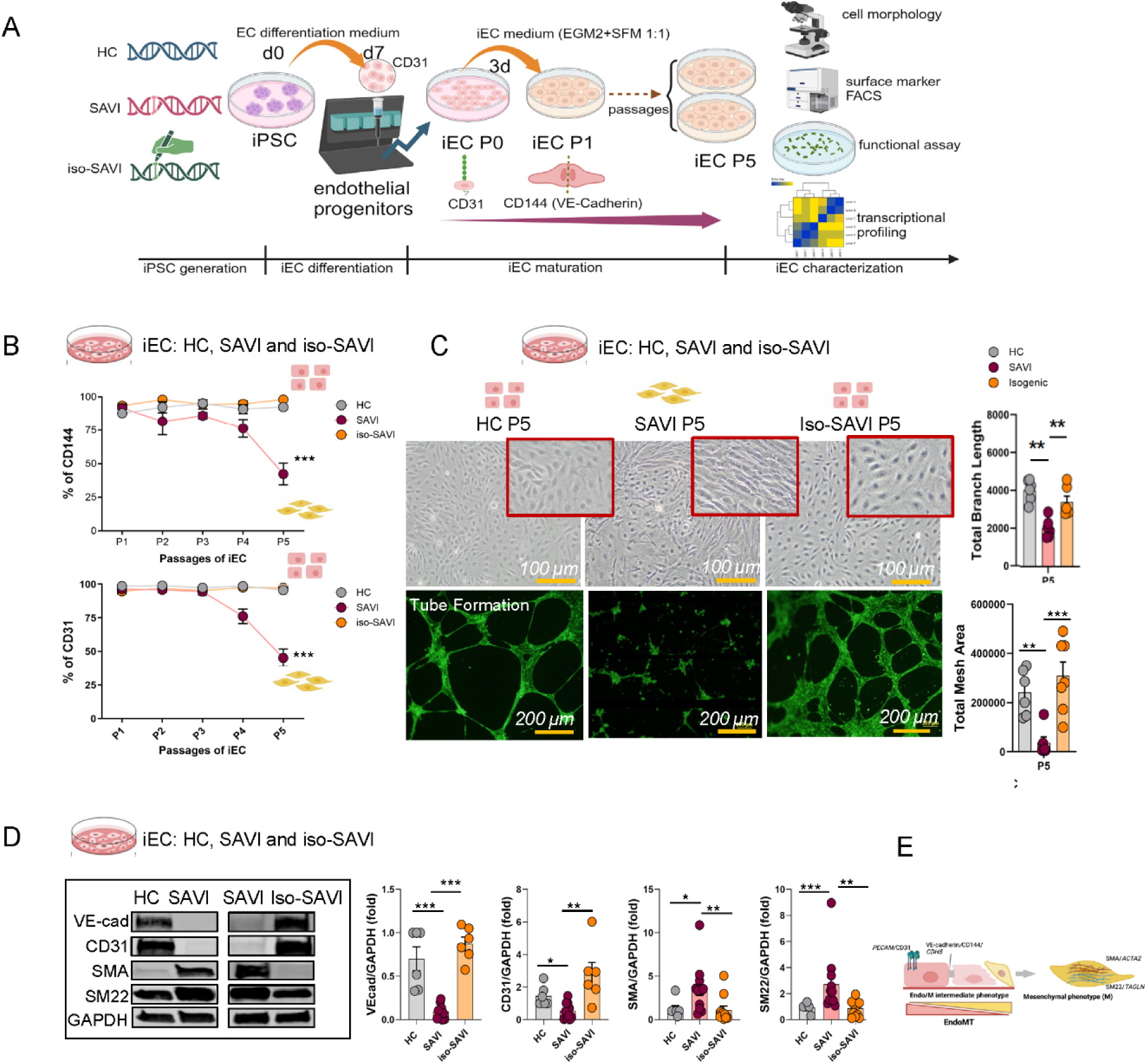
iPSC-derived endothelial cells (iECs) from SAVI patients but not from isogenic and HC iECs spontaneously undergo EndMT upon cell passage. **A.** Experimental workflow and group design (created with BioRender.com). **B.** Flow cytometric analysis of endothelial surface markers in HC, SAVI, and isogenic–SAVI (iso-SAVI) iECs. SAVI iECs progressively lost CD144 (VE-cadherin) and CD31 beginning at P3. Data summarize four HC- and SAVI-derived iEC lines and two iso-SAVI lines (mean ± SEM; ***p < 0.001, 2-way ANOVA). Representative flow cytometry profiles are shown in Figure S2C. **C.** Morphology and tube-formation capacity of iECs. Upper: HC and iso-SAVI iECs maintained cobblestone morphology from P1 to P5, whereas SAVI iECs transitioned to elongated fibroblast-like cells. Scale bar: 100 µm. Lower: Tube-formation assays performed at passages 1, 3, and 5; representative P5 images are shown (P1 images in Figure S2B). Total branch length and mesh area comparisons across groups are shown on the right (mean ± SEM; ***p < 0.001, **p < 0.01, *p < 0.05; 2-way ANOVA). Scale bar: 200 µm. **D.** Western blot analysis of endothelial (VE-cadherin, CD31) and mesenchymal (SMA, SM22) markers in iECs at P5. iECs were generated from three HC and three SAVI donors, with two iso-SAVI lines (three clones total). Representative blots and quantification (mean ± SEM; Mann–Whitney test) are shown. GAPDH served as a loading control. **E.** Schematic illustration summarizing panel D.

### Multi-Modal profiling of SAVI iECs identifies transcriptional drivers of STING-mediated EndMT

To explore the pathomechanisms driving EndMT in SAVI, we performed pathway enrichment analysis using differentially expressed genes (DEGs) from bulk RNA sequencing of SAVI and HC iECs and genetically corrected iso-SAVI lines. By P5, SAVI iECs displayed significant enrichment for pathways related to extracellular matrix organization, tissue fibrosis, wound healing and type I interferon (IFN) signaling (Figure 3A, Figure S3A-B). Targeted gene set analysis showed an EndMT signature and an IFN signature in SAVI iECs (Figure 3B, Figure S3C). An EndMT signature previously described in Human Umbilical Vein Endothelial Cell (HUVEC) and by endothelial lineage tracing in a murine model ^25^ confirmed a profound loss of an endothelial signature and gain of a mesenchymal signature in SAVI iECs at P5 (Figure 3B). Trends of these transcriptional changes were also observed in lesional alveolar regions from SAVI lung biopsy tissue (Figure S3D) and in human lung microvascular endothelial cells (HLMECs) treated with the STING agonist 2′3′-cGAMP for 5 days (Figure S3E). Cotreatment of HLMECs with cGAMP and IFNβ accelerated the cGAMP-induced loss of endothelial markers, however, IFNβ treatment alone largely preserved endothelial markers expressions and did not induce the upregulation of the mesenchymal signature. Along these lines blocking type1 IFN signaling did not preserve the endothelial markers/signature on cGAMP or cGAMP + IFNβ treated cells (Figure S3E) supporting the model in which cGAS-STING and not IFNβ is the primary driver of loss of the endothelial signature and induction of multiple mesenchymal genes with IFNβ acting as disease accelerator/amplifier.

**Fig3:**
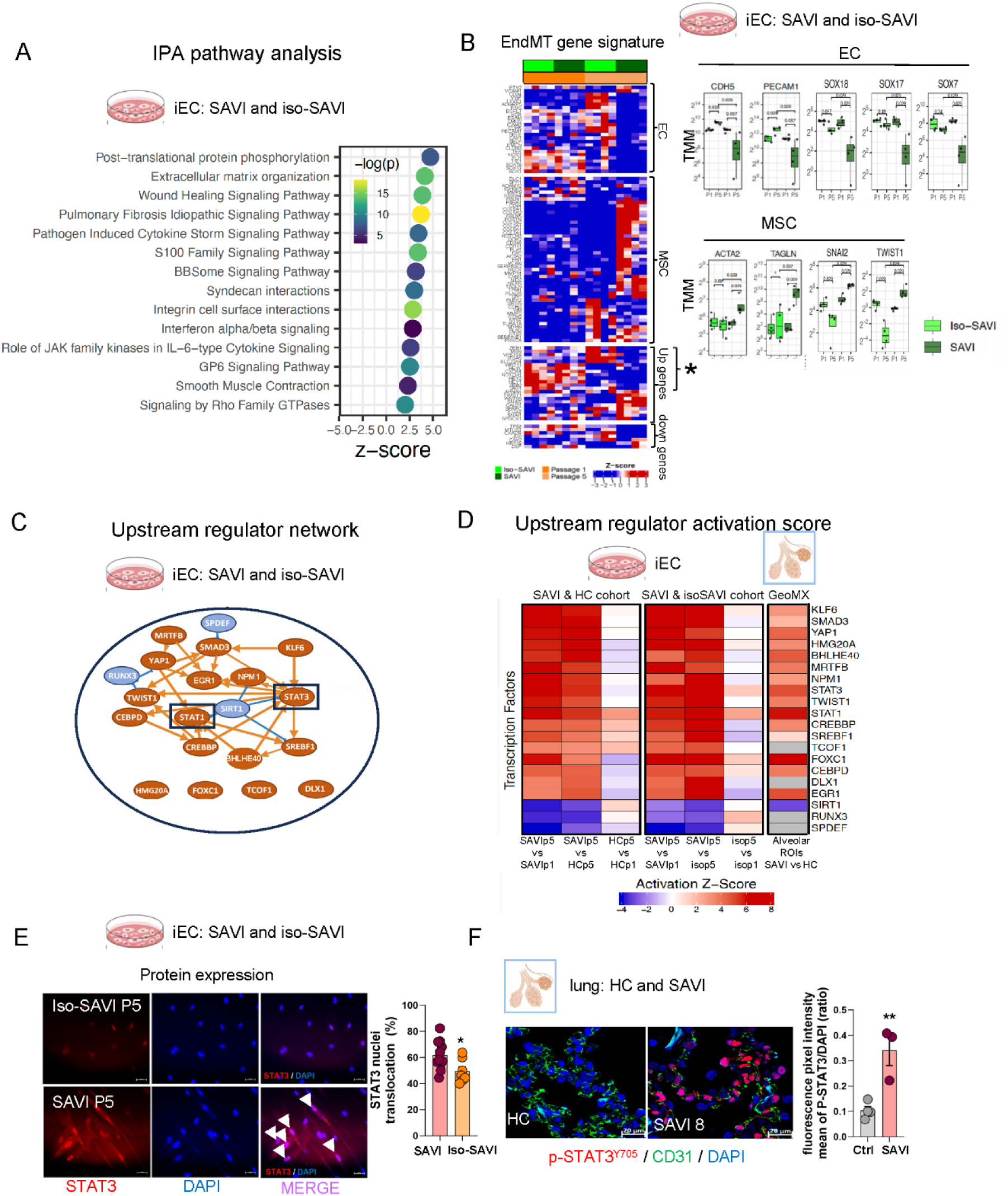
Multi-Modal profiling of SAVI iECs identifies transcriptional drivers of STING-mediated EndMT. **A.** Top pathways enriched in SAVI P5 vs. P1 (iso-SAVI_SAVI cohort) by Ingenuity Pathway Analysis (IPA). Positive z-scores indicate pathway activation. Additional HC vs. SAVI analysis is shown in Figure S3A–B. **B**. EndMT signature heatmap in parental SAVI and isogenic iECs, with box-and-whisker plots of representative endothelial, mesenchymal, and previously reported EndMT-associated genes. Asterisks mark genes upregulated under TGFβ-induced EndMT ^25^ but downregulated in SAVI iECs. **C.** IPA-derived transcription factor network of significant upstream regulators in the iso-SAVI_SAVI cohort (orange = activated; blue = inhibited). **D**. Heatmap of activation z-scores for the same transcription factors in panel C, showing similar patterns in SAVI vs. HC iECs and SAVI vs. HC lung tissue. **E**. Constitutive STAT3 nuclear translocation in SAVI iECs. STAT3 immunofluorescence (red) with DAPI (blue) at P5 from one SAVI and matched isogenic control line; white arrows indicate nuclear STAT3 (scale bar: 20 µm). Quantification reflects STAT3⁺ nuclei per total DAPI⁺ cells across 7–11 fields from 3–4 wells (mean ± SEM; *p < 0.05, two-tailed unpaired t-test). **F**. STAT3 activation in SAVI lung biopsies. Lung sections from 4 HC and 3 SAVI patients were stained for p-STAT3^Y705^ (red) and CD31 (green) with DAPI (blue). Representative images (scale bar: 20 µm) and quantification of p-STAT3 ^Y705^/DAPI ratios (≥5 images per patient) show increased STAT3 activation in SAVI (mean ± SEM; ***p < 0.001, two-tailed unpaired t-test).

In SAVI iECs, mesenchymal transcription factors (TF) *SNAI2*/SLUG and *TWIST1* were upregulated, while regulators commonly induced by Notch/Wnt or TGF-β signaling and implicated in cardiac cushion formation and fibrosis (*HEY1/2, HEYL, JAG1, NOTCH1, ZEB1/2*) ^26–29^ were downregulated; *SNAI1*/SNAIL, a TGF-β induced EndMT driver, remained unchanged (Figure 3B). These findings suggest a potential Notch/Wnt and TGF-β independent pathway involved in SAVI EndMT. Notably, the EC fate transcription factors belonging to the SOXF subgroup which include SOX18, SOX17 and SOX7 were all downregulated in SAVI iECs at P5. Only *SOX18*, a key transcription factor of endothelial identity and vascular development, was consistently downregulated across all models of STING activation including in patients’ lesional lung biopsies (Figure 3B Figure S3D and S3E). An upstream regulator analysis using Ingenuity Pathway Analysis (IPA) revealed a cooperative TF network centered on the inflammatory regulators *STAT3* and *STAT1*, with *STAT3* functioning as the principal node connecting most TFs in the network. Additional predicted activators included *SMAD3*, *TWIST1, FOXC1, YAP1,* factors commonly involved in TGF-β - and biomechanically mediated profibrotic gene regulation ^30–34^, as well as *CREBBP, CEBPD, NPM2.* Reduced *SIRT1* activity further suggests the involvement of epigenetic regulators that may converge in SAVI iECs ^35–38^ (Figure 3C). Similar upstream TFs were activated in SAVI lung tissues (Figure 3D). Consistent with these predictions, mRNA levels of *STAT1* and *TWIST1* were increased in SAVI iECs (Figure 3B, Figure S3F), while STAT3 transcription was unchanged. However, STAT3 protein was elevated and translocated to the nucleus in SAVI iECs at P5, and STAT3 phosphorylation was significantly increased in SAVI lesional lung biopsies compared with controls (Figure 3E-F), consistent with post-transcriptional activation.

In summary, multi-modal profiling identified loss of an endothelial program and induction of a mesenchymal signature that only partially overlapped with a published EndMT signature with upstream regulator analysis highlighting a role for STAT1/3 signaling and loss of the endothelial lineage transcription factors of the SOX family.

### Functional studies identify STAT3 as pivotal early driver of a SLUG dependent mesenchymal transcription program

Canonical cGAS/STING signaling induces STING/TBK1/IRF3/STAT1-mediated IFN responses^39,40^, which drive the upregulation of an IFN signature in whole blood of SAVI patients and serves as biomarker that correlates with systemic inflammation ^41^. In contrast, STAT3-mediated pathways were upregulated in SAVI IECs and in lesional lung tissue (Figure 3E-F), where they have previously been linked to mesenchymal transformation in EMT ^42,43^, suggesting activation of a non-canonical STING pathway in endothelial cells. To evaluate both pathways, we profiled STAT1 and STAT3 signaling in SAVI and control iECs, as well as in HLMECs and assessed their impact on mesenchymal transition and loss of endothelial markers during SAVI EndMT (Figure 4A). During serial passaging of SAVI iECs, phosphorylation of pSTAT1^Y701^ and pSTAT3^Y705^ peaked at P3; however, only pSTAT3^Y705^ remained significantly elevated at P5 compared to HC and iso-SAVI iECs, while STAT1^Y701^ phosphorylation declined to control levels (Figure 4B). To assess canonical signaling, we stimulated SAVI iECs with 2’3’-cGAMP at P2, prior to mesenchymal transition, and at P4, when cells had largely transitioned to a mesenchymal phenotype. At P2, cGAMP stimulation significantly increased pSTING^S366^, pTBK1^S172^ and modestly (but not significantly) increased STAT1^Y701^ phosphorylation in SAVI iECs compared to HC iECs. At P4, when SAVI iECs had largely transitioned to a predominantly mesenchymal phenotype, STING, TBK1 and STAT1 phosphorylation were similar to HC iECs whereas pSTAT3^Y705^ phosphorylation remained significantly elevated in SAVI iECs at both early (P2) and late (P4) passages (Figure S4A). Both cGAMP-induced acute STAT3 phosphorylation and constitutive STAT3 phosphorylation in SAVI iECs were blocked by STING (STINGi, IFM35883) or TBK1 (TBK1i, MRT67307) inhibitors, each reducing levels by ∼ 60% (Figure 4C-D), suggesting a STING/TBK1-mediated mechanism. Interestingly, cGAMP-induced IRF3 phosphorylation was transient and not significantly different in SAVI and HC iECs (Figure S4A). No constitutive activation of IRF3 was observed in SAVI iECs despite increased STAT3 activation (both, by phosphorylation and nuclear translocation respectively) (Figure 4B, Figure S4B), indicating that non-canonical STAT3 signaling in SAVI iECs is STING/TBK1-dependent but IRF3-independent.

**Fig4:**
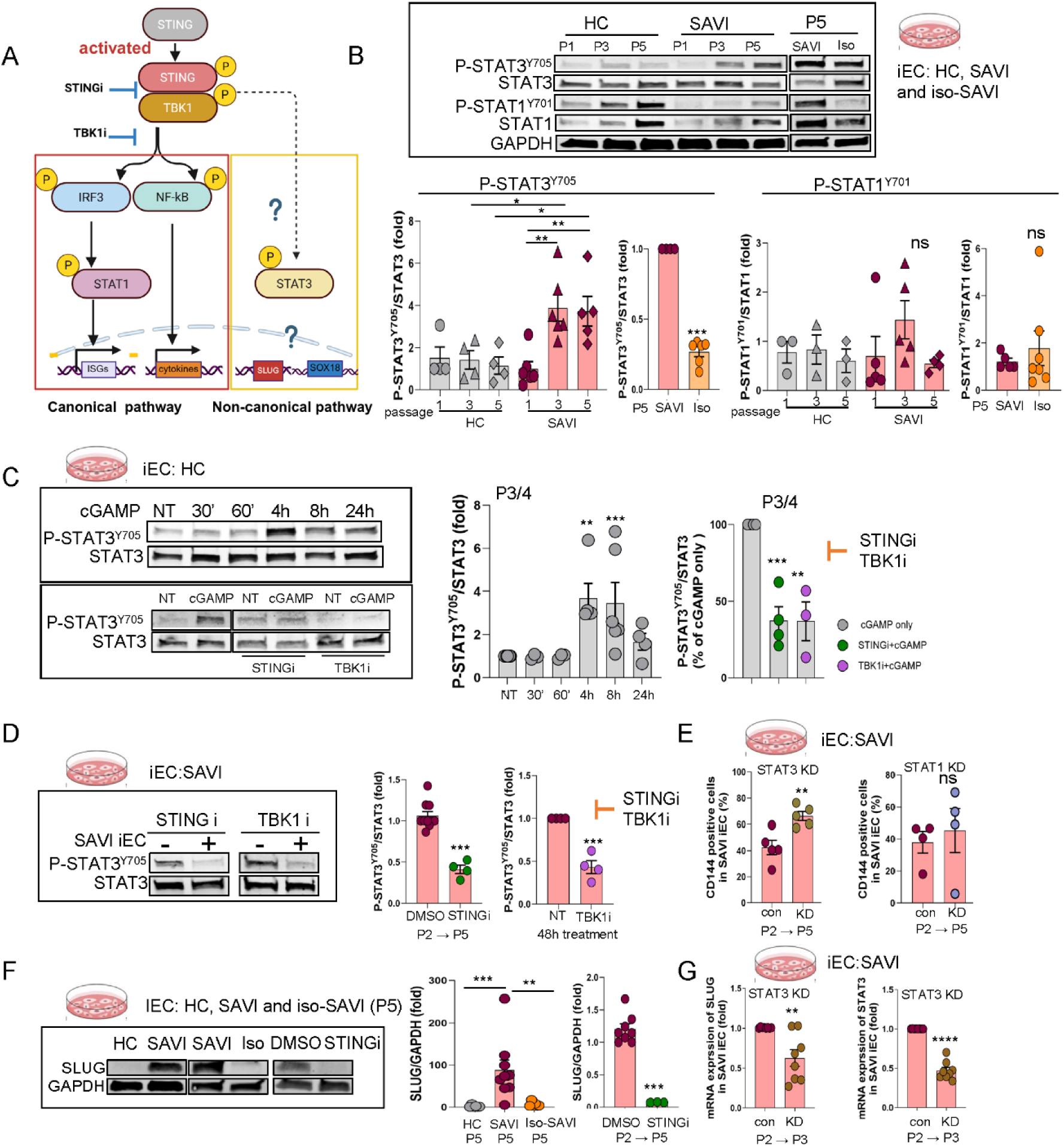
Functional studies identify STAT3 as pivotal early driver of a SLUG dependent mesenchymal transcription program. **A.** Diagram of the Canonical and Non-canonical STING signaling (created with BioRender.com). **B.** Constitutive STAT3 and STAT1 activation in SAVI iECs. Phospho-STAT3^Y705^ and phospho-STAT1^Y701^ (normalized to total protein) were measured in iECs from 3 HC and 3 SAVI donors across passages 1, 3, and 5. SAVI iECs showed increased baseline STAT3 activation (mean ± SEM; ***p < 0.001, **p < 0.01, *p < 0.05, unpaired t-test). **C.** STING/TBK1 inhibition blocks acute cGAMP-induced STAT3 activation in HC iECs. iECs from 3 HC donors were stimulated with 2’3’-cGAMP (20µg/ml) for 30 min–24 h. STING inhibitor IFM35883 (STINGi, 2.5 μM) and TBK1 inhibitor MRT67307 (TBK1i, 5 μM) were pre-treated one hour before cGAMP. STAT3 activation at 4h was prevented by STING/TBK1 inhibition (mean ±SEM, ***p < 0.001, **p < 0.01, Mann-Whitney test). **D.** STING/TBK1 inhibition reduces constitutive STAT3 activation in SAVI iECs. SAVI iECs were treated with IFM35883 (2.5 μM) from P2–P5 or with TBK1i (5 μM) for 48 h. Quantification is from four experiments in iEC line from patient SAVI1 (mean ±SEM, ***p < 0.001, unpaired t-test). **E.** STAT3 shRNA knockdown (KD), not STAT1 KD preserves CD144 (VE-cadherin) in SAVI iECs. iECs from 3 individual SAVI patients infected with control, STAT3, or STAT1 shRNA at P2 were analyzed by flow cytometry at P5 (mean ±SEM, **p < 0.01, paired t-test). Representative flow cytometry profiles are shown in Figure S4C. **F.** Constitutive protein expression of SLUG/*SNAI2* is elevated in SAVI iECs and suppressed by STING inhibition. Protein from 3 HC, 3 SAVI, and 3 iso-SAVI iECs at P5 showed increased SLUG (normalized by GAPDH) in SAVI; IFM35883 (2.5 μM) treatment from P2–P5 prevented SLUG upregulation in SAVI iECs (SAVI 1) (mean ±SEM, ***p < 0.001, **p < 0.01, unpaired t-test). See also Figure S4E. **G.** STAT3 shRNA knockdown reduces SLUG mRNA expression (normalized by internal control 18S) in SAVI iECs at P3. Data are summarized from 3 individual SAVI patients iEC lines. *p < 0.01 as determined by two-tailed unpaired t-test.

STAT3 knockdown, but not STAT1 knockdown, partially preserved the expression of endothelial marker VE-cadherin (CD144/*CHD5*) in SAVI iECs (42% expression with control shRNA vs 66% with STAT3 shRNA Figure 4E, Figure S4C), as well as in cGAMP+IFNβ-stimulated HLMECs (Figure S4D), highlighting a pivotal role of increased STAT3 activation in STING-induced EndMT. Since previous studies have shown that STAT3 selectively binds the *SNAI2*/SLUG promoter, but not *SNAI1*/SNAIL, *TWIST1*, *ZEB1* or *ZEB2,* promotors, and activates a mesenchymal transcriptional program ^44–46^, we tested the STAT3/SLUG axis. SLUG expression was significantly upregulated in SAVI iECs at P5 compared to HC iECs (Figure 4F, Figure S4E) and was suppressed by STING inhibition or STAT3 knockdown (Figure 4F-G) and similarly reduced in cGAMP-stimulated HLMECs treated with the STAT3 inhibitor Stattic (Figure S4F). These findings identify IRF3-independent STAT3 activation as a driver of a SLUG-dependent mesenchymal transcription program in SAVI endothelial cells.

### STING-driven enhancer remodeling targets GATA2/AP-1/SOX18 networks that maintain endothelial stability

While homeostatic STAT3 signaling is essential for endothelial homeostasis ^47^, chronic activation of the STING-STAT3 axis in SAVI iECs led to progressive loss of endothelial makers, prompting evaluation of chromatin changes that might drive repression of an endothelial maintenance program. ATAC-seq revealed a more than 3-fold increase in dynamic chromatin regions (CRs), in SAVI iECs compared with HC iECs by passage 3, largely driven by chromatin closing (44,064 vs. 13,513; *p<0.0001*) (Figure 5A-B, Figure S5A). Closing CRs were enriched for AP-1, ETS, and SOX motifs (Figure 5C, Figure S5B), consistent with reduced accessibility for endothelial lineage-stabilizing transcription factors ^48–50^.

**Fig5:**
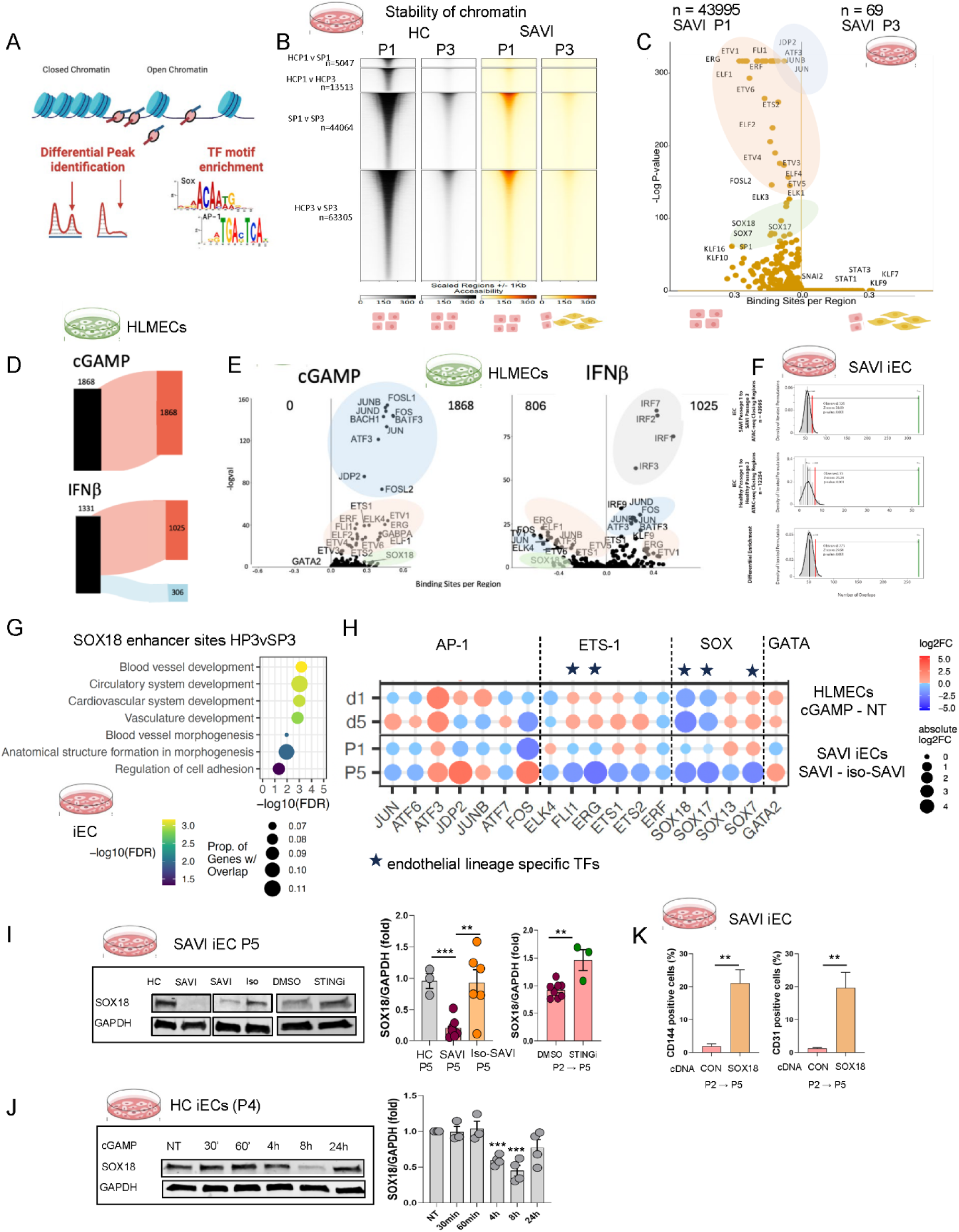
STING-driven enhancer remodeling targets GATA2/AP-1/SOX18 networks that maintain endothelial stability. **A.** Workflow showing ATAC-seq, motif enrichment and pathway analyses. **B**. Heatmap of differentially accessible regions in HC and SAVI iECs at P1 and P3 (ATAC-seq; chi-square with Yates correction). **C.** Motif enrichment in regions differentially accessible between SAVI P1 and SAVI P3. Motifs enriched in the 43,995 SAVI P1–open regions appear on the left; motifs enriched in the 69 SAVI P3–open regions appear on the right. Additional comparisons are in Figure S5B. **D.** ATAC-seq in HLMECs stimulated with 2’3’-cGAMP or IFNβ for 8 h. Sankey plot shows the number of gained and lost accessible regions relative to non-treated (NT) cells. **E.** The enrichment of transcription factor binding motifs within the differentially accessible regions between the non-stimulated (NT), 2’3’-cGAMP or IFNβ stimulated HLMEC ATAC-seq libraries. Motif enrichment in regions more accessible after cGAMP stimulation in HLMECs (1,868 regions; none were more open in NT cells). Motif enrichment within the 1025 regions with increased accessibility in the IFNβ stimulated condition is shown on the right side of the y = 0 line. Cloud color as in B grey cloud indicates IRF TF motifs that are becoming increasingly accessible. **F.** These plots show the results of permutation tests from the regioneR package for the overlap between SOX18 ChIP-seq peaks from the Overman et al dataset and the SAVI P1 to SAVI P3 closing regions, or the HC P1 to HC P3 closing regions. A differential permutation test was also conducted to compare the relative enrichment of SOX18 ChIP-seq peaks in these two sets of regions. The green bar indicates the observed number of overlaps between the datasets. The black bar indicates the mean value of overlaps between the transcription factor ChIP-seq peaks and the random permuted regions. the distribution of overlaps from the permutations are shown as the gray histogram). The red bar indicates the number of overlaps at the threshold of significance p = 0.05. Differential enrichment analysis (bottom plot) showed significant overrepresentation of SOX18 binding sites in SAVI closing CRs at P3 (273 regions, z = 29.54 ; p < 0.001). **G**. The pathway enrichment of genes annotated by GREAT to differentially accessible regions from the SAVI iEC P1 to SAVI iEC P3 conditions that exhibited a decrease in accessibility in the P3 cells. These regions (and annotated genes) were filtered by the regions that overlapped SOX18, JUN, FOS, and GATA2 ChIP-seq peaks from the HUVEC ChIP-seq data sets listed in the methods section. **H.** Gene expression changes of endothelial lineage transcription factors across SAVI vs. iso-SAVI iECs (P1, P5) and cGAMP-treated vs. NT HLMECs (d1, d5) by RNAseq analysis. Red points indicate higher expression in SAVI or cGAMP conditions; endothelial TFs are starred. **I.** Reduced SOX18 protein in SAVI iECs at P5 compared with HC and iso-SAVI lines (3 individual iEC lines for each group), and partial restoration (∼50%) following STING inhibitor treatment (IFM35883, 2.5 μM, P2–P5) in SAVI iECs (SAVI1, n = 3). GAPDH served as loading control. Data are represented as mean ± SEM; ***p < 0.001, **p < 0.01, two-tailed unpaired t-test. **J.** SOX18 downregulation in HC iECs following 20 μg/ml 2’3’-cGAMP stimulation (30 min–24 h), peaking at 8 h (∼50% reduction). Quantification from three HC donors (mean ±SEM, ***p < 0.001, two-tailed unpaired t-test). **K.** SOX18 overexpression preserves endothelial surface markers CD144 and CD31 in SAVI iECs at P5. SAVI iECs were transduced at P2 with control or SOX18 cDNA; Flow cytometry analysis at P5 shows increased CD144⁺ and CD31⁺cells in SOX18 overexpression group. Data are represented as mean ± SEM; ***p < 0.001, **p < 0.01, two-tailed unpaired t-test. (n = 4, SAVI1 iEC line).

To distinguish cGAMP/STING-specific from IFNβ- driven effects and to exclude potential reprogramming artifacts in the iECs, which exhibit greater plastic capacity than mature ECs ^51,52^, we profiled primary HLMECs following short-term 2’3’-cGAMP-mediated STING activation or IFNβ stimulation for 8 hours. Short-term cGAMP stimulation induced 1,868 newly accessible chromatin regions enriched for AP-1 and ETS motifs, whereas IFNβ induced fewer regions dominated by IRF motifs (Figure 5D-E) ^53^. Notably, 85.3% of regions that became accessible following acute (8h) cGAMP stimulation (n=1593) overlapped with chromatin loci that became inaccessible under chronic STING activation in SAVI iECs, compared with only 5.2% of IFNβ–induced regions (Figure S5C, left panel). Pathway analysis of genes associated with these regions revealed enrichment for pathways related to endothelial integrity and vascular homeostasis (Figure S5C, right panel), suggesting that these CRs may be regulated by cGAMP-responsive endothelial enhancers that become progressively repressed under chronic STING signaling.

Because BRD4 re-localization serves as key integrator of inflammatory chromatin remodeling, we next examined whether BRD4, a BET-family chromatin reader that does not bind DNA directly but occupies acetylated active enhancers and is known to redistribute from homeostatic endothelial enhancers to stress-responsive regulatory elements,^54,55^ is enriched at cGAMP-induced changing chromatin regions. We therefore performed enrichment analyses integrating public ChIP-seq datasets for BRD4, and from TFs with well-established roles in regulating vascular identity ^56^ including the ETS-family regulator, ERG,^57^ as well as SOX18, GATA2, ^58,59^ and the AP-1 complex (JUN/FOS) ^55^ to validate the motif enrichment analysis. Chromatin regions rendered accessible by acute cGAMP stimulation (1,593 cGAMP-responsive regions) as well as regions that progressively lost accessibility during SAVI iEC passaging (P1 to P3) were significantly enriched for BRD4- and AP-1–associated peaks, consistent with the engagement of an inflammatory BRD4 program; SOX18 and ERG peaks were also enriched (Figure S5D).

Gene expression changes previously associated with acute BRD4 re-localization ^54,55^ were recapitulated in both cGAMP-stimulated endothelial cells and in SAVI iECs, including repression of *SOX18* and *EGFL7* ^54^, two of the most strongly downregulated endothelial genes in SAVI lung tissue, as well as additional genes involved in endothelial stability (Figure S3D, Figure S5E). The closing CRs in SAVI iECs were significantly more enriched for SOX18 ChIP-seq peaks compared to HC iECs . Of 326 SOX18 ChIP-seq peaks that lost accessibility in SAVI iECs at P3 (z = 38.0, p < 0.001), 53 were significantly enriched in HC iECs at P3 (z = 25.24, p < 0.001). Differential enrichment further identified 273 SOX18 peaks within SAVI-closing regions (z = 29.54, p < 0.001), indicating a strong, disease-specific loss of SOX18-associated accessibility (Figure 5F). Similarly, ERG ChIP-seq peaks were also enriched in closing CRs in SAVI iECs (Figure S5F). SOX18 ChIP-seq peaks were more co-localizing with AP-1 and/or GATA2 compared with ERG, the latter showing significantly less colocalization with AP-1 and/or GATA2, (Yates-corrected χ² p < 0.0001) (Figures S5G). This pattern is consistent with a role for SOX18 as a stabilizing enhancer at regulatory elements controlling critical vascular development pathways (Figure 5G). and downstream target genes including *HHEX, MECOM, TIE1,* and *CALCRL*) (Figures S5H), all of which were transcriptionally downregulated in mostly EMT transitioned SAVI iECs compared to HC iECs that maintained their endothelial phenotype at P5.

Together these data indicate that acute STING activation, but not IFNβ signaling, perturbs the endothelial enhancer landscape likely by redirecting BRD4 to AP-1-enriched inflammatory regulatory elements, leading to transcriptional repression of SOX18 ^54^. Consistent with this model transcriptional profiling revealed early repression of *SOX18* and *SOX17* following acute cGAMP stimulation, whereas broader suppression of endothelial transcription factors emerged only under chronic STNG activation in SAVI iECs (P5). ETS-family TFs (*ERG, FLI1, ETS1/2*) were transiently induced after acute cGAMP exposure but were consistently suppressed during chronic STING activation in SAVI iECs (P5) (Figure 5H Figure S5I). Notably, HDAC9, a class IIa deacetylase implicated in EndMT and vascular remodeling ^49,60^ was induced early following cGAMP stimulation (Figure S5I), suggesting early engagement of enhancer-modifying mechanisms during acute inflammatory signaling.

At the protein level, low SOX18, an endothelial transcription factor with a nonredundant role in endothelial differentiation and unique homodimerization capacity among SOX family members, ^49,61^ was markedly reduced in SAVI iECs (P5) and increased by 54% following sustained STING inhibition (IFM35883) (Figure 5I). In cGAMP-stimulated HC iECs, SOX18 protein expression nadired at 8 hours and was restored by inhibition of STING (IFM35883), TBK1 (MRT67307), or STAT3 (Stattic) (Figure 5J, Figure S5J). In contrast, repression persisted beyond 24 hours despite STAT3 blockade (Figure S5K). Functionally, SOX18 overexpression partially preserved the endothelial surface markers CD144 (VE-cadherin) and CD31 (Figure 5K) and induced upregulation of the key endothelial transcription factor ERG, with ERG mRNA expression increased by ∼20 fold compared with control SAVI iECs at P5 (Figure S5L).

Collectively, these findings support a model in which acute STING-driven chromatin remodeling follows a pattern of inflammatory BRD4 redistribution associated with SOX18 suppression ^54^, whereas chronic STING activation leads to selective collapse of SOX18-enriched endothelial maintenance enhancers. This process coincides with progressive destabilization of GATA/ETS/SOX/AP1-enriched chromatin regions ^62–64^ that sustain an endothelial maintenance network, resulting in loss of endothelial identity that can be partially restored by STING inhibition or SOX18 re-expression.

### STING inhibition but not drugs approved for treatment of pulmonary fibrosis preserve the endothelial phenotype in STING-induced EndMT

Having identified that STING activation drives chromatin remodeling and transcriptional changes associated with mesenchymal transition, we next sought to investigate whether the STING-induced chromatin and transcriptional reprogramming that drives mesenchymal transition contributes to fibrotic signaling in SAVI. Notably, the EndMT transcriptional program observed in SAVI iECs was also enriched in alveolar tissue from SAVI patients, suggesting a link between STING activation, endothelial dysfunction, and lung fibrosis. To evaluate whether targeting downstream signaling pathways could preserve endothelial identity, we examined the effects of JAK1/2 inhibition in cGAMP-stimulated HLMECs and in spontaneously transitioning SAVI iECs. Baricitinib, a reversible JAK1/2 inhibitor used clinically for SAVI ^41,65^, partially restored expression of VE-Cadherin (CD144)/*CDH5* and CD31/*PECAM1* and suppressed upregulation of mesenchymal marker SMA/*ACTA2* in the HLMECs model after 5 days of stimulation with 2’,3’ cGAMP +/- recombinant IFNβ. (Figure S6A-C). However, in SAVI iECs, baricitinib had a limited protective effect. Co-culture with baricitinib from passage 2 to passage 4/5 preserved VE-Cadherin (CD144)/*CDH5* but not CD31/*PECAM1* and did not prevent upregulation of SMA/*ACTA2* at later passages (Figure 6A-B).

**Fig6:**
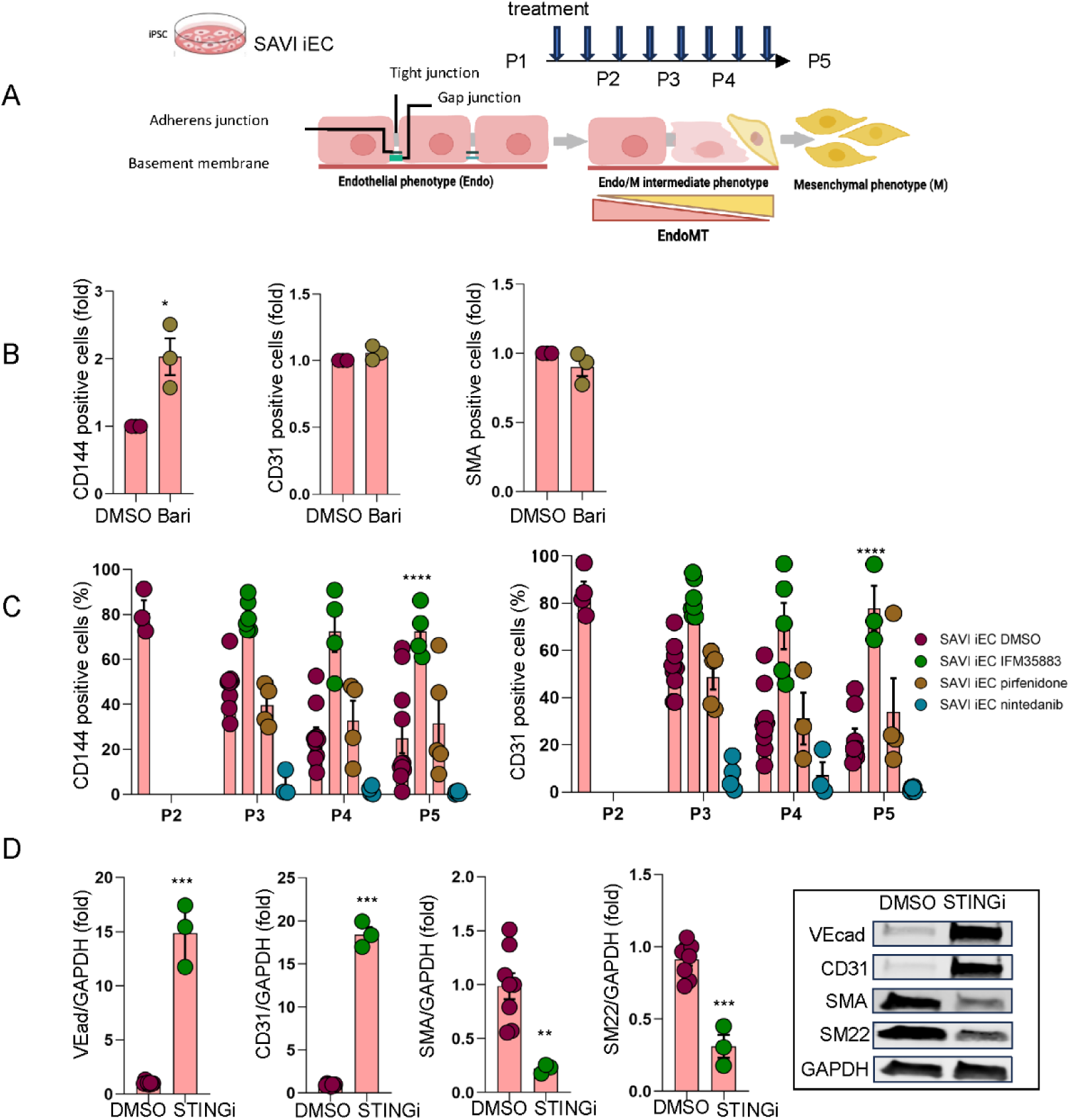
STING inhibition but not drugs approved for treatment of pulmonary fibrosis preserve the endothelial phenotype in STING induced EndMT. **A.** Schematic of drug testing in the SAVI in-vitro disease model. **B.** Partial rescue of SAVI iEC EndMT by the FDA-approved JAK/STAT inhibitor Baricitinib. SAVI iECs were treated with Baricitinib (1 μM) from P2 to P3–4 (early harvest due to toxicity). Flow cytometry analysis of CD144 (VE-cadherin) and CD31 showed modest preservation of endothelial markers versus DMSO. Data from three SAVI donors (mean ± SEM; *p < 0.05; two-tailed t-test). Effects in cGAMP-treated HLMECs are shown in Fig. S6A–C. **C.** Impact of STING inhibition and FDA-approved antifibrotic drugs on EC surface markers. SAVI iECs were treated from P2 with IFM35883 (2.5 μM), pirfenidone (10 μM), nintedanib (1 μM), or DMSO. STING inhibition nearly fully preserved CD144 (VE-cadherin) and CD31, pirfenidone had no effect, and nintedanib markedly worsened EndMT. Representative flow cytometry profiles in Fig. S6E. Data from ≥3 experiments in the SAVI1 line (mean ± SEM; *****p < 0.0001; 2-way ANOVA). **D.** STING inhibition normalizes EndMT markers in SAVI iECs. SAVI iECs treated with IFM35883 (2.5 μM) or DMSO from P2–P5 were analyzed by Western blot. VE-cadherin and CD31 increased, while SMA and SM22 decreased with STING inhibition. Quantification from three experiments in SAVI1; representative blots shown (mean ± SEM; ***p < 0.001, **p < 0.01; two-tailed unpaired t-test).

To further probe mechanisms of fibrosis, we tested two FDA-approved anti-fibrotic agents, pirfenidone and nintedanib, in the SAVI-specific iEC model. Pirfenidone is known to reduce collagen synthesis and growth factor production ^66,67^, while nintedanib is a tyrosine kinase inhibitor (TKI) that targets multiple tyrosine kinases involved in fibroblast proliferation ^68^. Treatment began at P2, when over 80% of cells expressed endothelial markers (i.e. CD144 (VEcad) and CD31). Unlike pirfenidone or nintedanib, the STINGi (IFM35883) almost completely prevented spontaneous EndMT in SAVI iECs (Figure 6C, Figure S6D). Nintedanib unexpectedly accelerated loss of endothelial markers without affecting viability (Figure S6D-E), whereas pirfenidone had no measurable effect. STINGi (IFM35883), in contrast, preserved VE-cadherin and CD31 expression and suppressed protein expression of mesenchymal markers, SMA and SM22, at P5 (Figure 6D). These findings support a central role for STING in driving mesenchymal transition and suggest that the SAVI iEC and cGAMP-stimulated HLMEC models provide useful systems for evaluating candidate therapies targeting EndMT in inflammatory lung fibrosis.

## DISCUSSION

Our study identifies cGAS/STING as a driver of STAT3/SLUG-dependent, TGFβ-independent EndMT and as a repressor of SOX18, a core regulator of endothelial maintenance. The spontaneous EndMT observed in SAVI iECs identifies STING activation as a cell-intrinsic driver of endothelial plasticity. Inhibition of STING, TBK1, or to a lesser extend STAT3, mitigated EndMT, linking this signaling axis to the loss of endothelial identity and acquisition of a mesenchymal phenotype. EndMT occurred independently of IRF3, diverging from canonical STING-IFN signaling. Notably, cells during the transition from an endothelial to a mesenchymal-like phenotype retained responsiveness to cGAMP–IRF3 activation and continued to express IFN-stimulated genes, suggesting that IRF3-dependent signaling maintains immune responsiveness, while IRF3-independent STING–STAT3 signaling drives mesenchymal reprogramming and fibrotic remodeling.

While EndMT is essential during embryogenesis, for example, in the formation of cardiac valves and septa ^69^, it’s aberrant activation drives fibrotic remodeling in organs such as the lung, heart, and kidney ^70^. The wide range of cell types capable of undergoing mesenchymal transition underscores the need for disease-specific approaches to define the initiating cell populations and contextual cues involved. In idiopathic pulmonary fibrosis (IPF), alveolar epithelial injury and fibroblastic foci, characterized as clusters of proliferating fibroblasts and myofibroblasts embedded within extracellular matrix, represent key sites of active fibrogenesis and hallmark pathological features ^71–73^. In contrast, fibroblastic foci are absent in SAVI lung tissue ^74,75^. Consistent with these findings, SAVI fibroblasts produce type I IFN in response to cGAMP/STING activation but fail to adopt a myofibroblast phenotype, indicating that STING does not directly drive fibroblast-to-myofibroblast transition. Instead, SAVI lung pathology is dominated by pronounced endothelial injury, venulitis, smooth muscle actin (SMA) deposition, and marked remodeling in alveolar septae. Similar features of vascular remodeling can be seen in pulmonary venous obstruction and congestive heart failure ^76^ suggesting a model in which endothelial cells, rather than fibroblasts, serve as primary initiators of fibrosis. Together with the EndMT signature in SAVI lung tissue and murine studies implicating endothelial dysfunction in disease pathogenesis ^77^, our findings link *STING1* GOF mutations to spontaneous EndMT, identifying endothelial cells as key targets of STING hyperactivation and major drivers of fibrotic remodeling.

STING GOF mutations in SAVI iECs induce sustained STAT3 phosphorylation and upregulation of SLUG/*SNAI2*, and TWIST/*TWIST1*, which are key TFs shared by both epithelial-to-mesenchymal transition (EMT) and EndMT ^25,28^. Although STAT3 is well-established in promoting EMT in cancer ^43^, its activation downstream of STING in endothelial cells represents a previously unrecognized mechanism of EndMT initiation. The STING-STAT3-SLUG axis operates independently of TGFβ and diverges from classical developmental and cytokine-induced EndMT pathways, as TFs governing cardiac EndMT and TGFβ-induced transitions such as SNAIL/*SNAI1, NOTCH1, JAG1, HEY1/2* ^27^ and *ZEB1/2* ^78,79^, were not induced and were in some cases suppressed. Thus STING emerges as a noncanonical initiator of mesenchymal reprogramming that converges on a transcriptional programs broadly conserved across EMT and EndMT, yet distinct in its upstream drivers ^28^.

Epigenetic profiling revealed extensive chromatin remodeling in SAVI iECs and cGAMP-treated HLMECs that was not recapitulated by IFN-β, underscoring the specificity of STING-signaling. Chromatin regions that transiently opened upon acute STING activation became selectively inaccessible in chronically activated SAVI iECs, suggesting that persistent signaling converts transient enhancer activation into stable repression. These sites were enriched for SOX18-and ERG-associated enhancer elements, consistent with loss of lineage-stabilizing TF activity. SOX18 emerged as one of the earliest and most consistently repressed TFs in both systems, coinciding with reduced accessibility at SOX18-bound endothelial maintenance loci including the TFs *HHEX* ^80,81^, and *MECOM* ^82^, and vascular stability genes *CALCRL*, *TIE1, SLC12A2* and *RASGRP3* ^83^.

Early repression of SOX18 may reflect acute STING-driven redistribution of BRD4 from lineage-stabilizing to inflammatory regulatory elements, paralleling BRD4 dynamics in TNF-activated endothelial cells ^54^. Sustained STAT3 activation may further stabilize this chromatin state by maintaining NFκB signaling ^84^ and reinforcing BRD4 occupancy at stress-responsive loci. Together, these data suggest that acute STING activation imposes an initial epigenetic insult that weakens SOX18-dependent enhancer stability. In parallel with BRD4 redistribution, cGAMP stimulation transiently induced HDAC9, a class IIa histone deacetylase that remodels enhancer architecture primarily through protein-protein interactions rather than direct DNA binding ^85,86^. HDAC9 has been implicated in EndMT-associated vascular pathology, including atherosclerosis in murine models ^60^ and its selective induction downstream of STING suggests an early role in enforcing enhancer deacetylation and destabilization of endothelial enhancer programs. Although we did not assess direct interaction or colocalization between HDAC9 and SOX18, the coordinated engagement of HDAC9 induction and BRD4 redistribution indicates that multiple epigenetic modifiers are recruited early during STING activation, creating a chromatin environment in which chronic STING activation progressively converts transient inflammatory activation into stable chromatin repression, thereby destabilizing a SOX18-regulated endothelial maintenance program and locking endothelial cells into a mesenchymal state.

Consistent with this model, SOX18-dependent enhancer modules appear particularly sensitive to inflammatory remodeling. Partial recovery of endothelial markers upon SOX18 overexpression supports a stabilizing role for SOX18 at AP-1/GATA2 co-bound regulatory sites essential for endothelial identity ^49,59,87^. Notably, SOX18 ChIP-seq peaks co-localize with H3K4me1 and H3K27ac marks at intragenic regions of ERG and FLI1 (Figure S5M-N), consistent with SOX18 binding at active lineage-maintenance enhancers embedded within these endothelial TF genes. Such enhancers frequently function as constituent elements of broader regulatory domains, including super-enhancers, that sustain endothelial identity. Analogous to SOX9 loss during vascular development ^88^, loss of SOX18 may destabilize GATA2/AP-1/BRD4 containing regulatory complexes and promote TF redistribution toward mesenchymal reprogramming.

The convergence of transcriptional and epigenetic programs downstream of STING has important therapeutic implications. Current FDA-approved antifibrotics pirfenidone and nintedanib failed to preserve endothelial markers in vitro; nintedanib even exacerbated their loss, indicating that these agents do not effectively counteract the endothelial reprogramming central to SAVI-associated fibrosis. By contrast, inhibition of STING, TBK1, or STAT3, significantly attenuated mesenchymal transition and preserved endothelial identity, highlighting the STING–TBK1–STAT3 axis as a promising therapeutic target in inflammatory lung diseases. Endothelial integrity is maintained by a cooperative transcriptional–epigenetic network involving SOX18, GATA2, AP-1, and chromatin remodelers such as BRD4, HDAC9, and SWI/SNF (BAF) complexes ^89,90^. Rather than a single dominant factor, the cumulative weakening of multiple regulatory nodes likely drives irreversible transitions such as EndMT. Therapeutic strategies that reinforce the network at multiple levels may therefore be required to preserve epigenetic integrity and prevent maladaptive cell-fate transitions.

In summary, our study identifies a noncanonical STING-STAT3-SLUG axis as a key driver of TGFβ-independent EndMT and implicates repression of SOX18/AP-1/GATA2–dependent enhancer networks as central to endothelial dysfunction in SAVI. These findings clarify the molecular basis of vascular injury in STING-driven disease and highlight STING pathway inhibition as a compelling therapeutic strategy across fibrotic and inflammatory vasculopathies.

### Limitations of the Study

This study has several limitations. Sample size was constrained by the rarity of the disorder, limiting age- and inflammation-matched comparisons across disease stages. Although we integrated multi-omic profiling with functional assays, some mechanistic links remain correlative, and in vitro endothelial models may not fully reflect the in vivo vascular environment under chronic STING activation. The lack of longitudinal sample acquisition further limits resolutions of temporal dynamics, precluding definitive separation of early pathogenic drivers from downstream secondary effects. In addition, therapeutic perturbations were evaluated in vitro, and their efficacy and safety in vivo remain to be established. Despite these limitations, the convergence of transcriptional, epigenetic, and functional data provides a coherent framework for STING-mediated endothelial dysfunction and highlights candidate pathways and targets for future therapeutic investigation.

## METHOD

### Subjects and study approval

iPSCs were generated from fibroblasts derived from skin punch biopsy samples obtained from three SAVI patients (SAVI 1 (female, 10y), SAVI 2 (female, 15y), and SAVI 4 (male, 12y)) and from peripheral blood mononuclear cells (PBMNC) obtained from one SAVI patient (SAVI 3 (male, 26y)). Three healthy volunteers iPSCs lines were generated from skin fibroblasts (one male (age: 23 years old) and two females (ages: 66, 25 years old). iPSCs were generated previously by Dr. Bohem’s lab and NHLBI iPSC core ^91,92^. This study was approved by the NIAID’s institutional review board, and samples were collected under clinical protocol number 17-I-0016 (NCT02974595). Healthy volunteer samples were collected under clinical protocol number 10-H-0126 (NCT01143454) approved by the NHLBI’s institutional review board. All study participants patients provided written informed consent before participation per the ICH E6 Guidelines for Good Clinical Practice originating from the Declaration of Helsinki.

### Study participants

All patients were enrolled in natural history study NIH Protocol Number: 17-I-0016 (NCT02974595): Studies of the Natural History, Pathogenesis, and Outcome of Autoinflammatory Diseases (NOMID/CAPS, DIRA, CANDLE, SAVI, NLRC4-MAS, Still’s-like Diseases, and other Undifferentiated Autoinflammatory Diseases).

### Patient lung CT imaging evaluation

Features of lung inflammation and damage on Computed Tomography (CT) were clinically evaluated by a single radiologist (LF). Severe lung disease was defined as presence of at least one characteristic: lung fibrosis, respiratory insufficiency, pulmonary hypertension, or digital clubbing associated with interstitial lung disease on chest CT.

### H&E, Masson-Trichome, and Pathology scoring evaluation

Semi-quantitative scoring was based on the results of immunohistochemistry that assessed the inflammatory and chronic damage findings in lung biopsy tissues in 7 patients with SAVI and 5 anonymized control subjects with reported normal lung biopsy. The inflammatory score was composed of inflammatory cell infiltrate including alveolar macrophages, interstitial histiocytes, neutrophils, lymphocytes, and intraparenchymal or peribronchial lymphoid aggregates. The chronic damage score was composed of fibrosis (perivascular, large and medium size vessels or perivascular, alveolar septum and interstitium), type 2 pneumocyte hyperplasia, hyaline matrix/membranes, cholesterol clefts, mucus plugging, transudate/exudate, emphysema (severity and percentage of lung parenchyma involvement), consolidation (percentage of lung parenchyma involvement), vascular damage/vasculitis, neovascularization, and thrombosis. Each aforementioned feature was graded 0, absent or within normal limits; 1, barely perceptible; 2, obvious but not striking; 3, striking. SAVI patients’ scores were compared with controls’ scores by Mann-Whitney test.

### Histology scoring of IPF and SAVI samples of H&E samples

Formalin-fixed, paraffin-embedded lung sections were reviewed and scored using a semi-quantitative ordinal scale. For each specimen, the following features were evaluated: pleural fibrosis, pleuritis, septal fibrosis, septal proliferation, follicular bronchiolitis, alveolar wall thickening, and fibroblast foci. Each feature was assigned a severity score from 0 to 3 (0 = absent/none, 1 = mild, 2 = moderate, 3 = severe) based on overall extent and intensity observed across the section at low and high magnification. Scoring was performed in a blinded manner by one experienced pathologist (HM) with respect to diagnosis (SAVI vs IPF). One representative section was assessed per case.

### Tissue immunofluorescence staining

Formalin-fixed, paraffin-embedded lung samples were deparaffinized with two 30-minute washes in xylene, followed by a series of two 100% ethanol washes for 3 minutes each, and then 3-minute washes in 90%, 80%, 70%, and 50% ethanol, concluding with water. Antigen retrieval was performed in a solution of 10 mM sodium citrate (pH 6.0) for 40 minutes at 95°C. Once cooled to room temperature, samples were blocked for one hour in 5% BSA, 20% donkey serum, and 0.1% Triton-X100 in PBS. Primary antibodies for CD31 (1:50, DAKO Cat#M0823), VE-cadherin (VEcad) (1:100, Santa Cruz Cat# sc-9989), or Phosphorylated STAT3 (P-STAT3^Y705^) (1:100 Cell Signaling Technology Cat#9145S) were applied overnight at 4°C. This was followed by the corresponding fluorescence-conjugated secondary antibody 1:1000 (Invitrogen) for one hour at RT (room temperature). FITC-conjugated SMA (1:200, Sigma-Aldrich F3777) was applied with second antibody when co-staining with VE-cadherin. Slides were mounted using DAPI-containing mounting media (Vector Laboratories) and imaged using a Zeiss LSM 880 confocal microscope. Fluorescence images were processed and further evaluated using the ZEN (blue edition) software.

### CODEX (Phenocycler Fusion)

For the Phenocycler Fusion multiplex protein detection assay, a slide with multiple lung FFPE sections from the SAVI patient (5 um thick tissues) were processed and stained with an antibody cocktail containing a mix of commercially available and custom conjugated antibodies (45) following recommendations from Akoya Biosciences.

The slide was baked overnight at 60°C to improve attachment of tissues to glass. It was immersed in xylene (2 times 10 min) to remove paraffin and later rinsed in a decreasing alcohol series to rehydrate the tissues (100%, 100%, 90%, 70%, 50%, 30% ethanol in MilliQ water for 4 min each). Antigen retrieval was performed in a pressure cooker set to low pressure using antigen retrieval solution pH 9 for 15 min. After the slides cooled down to RT, they were soaked in Hydration (2 times 2 min) and Staining buffer (up to 20 min). Antibodies were combined in staining buffer containing J, N, S and G blocking reagents and were added to tissues overnight at 4oC. The slide was washed in staining buffer (2 times 2 min) and the bound antibodies were fixed using 1.6% paraformaldehyde (diluted in storage buffer) for 10 min. The slide was washed in PBS (3 times 2 min), incubated in ice cold methanol for 5 min and, after additional PBS washes (3 times 2 min), were incubated in the crosslinker solution diluted in PBS for 20 min. After additional PBS washes (3 times 2 min) the stained slide was imaged on Phenocycler fusion. For imaging, a flowcell was created on the slide and the imaging run was set-up following the vendor’s recommendations. Details of antibody clone information are included in Key Resource Table.

At the end of the run, the generated 8-bit images were further processed to generate 16-bit images to reduce signal saturation on selected channels. Both the 8 and 16-bit images were uploaded into HALO image analysis software (NCI-HALO version 3.6) for visualization and analysis.

#### Antibody-conjugation with oligonucleotide-tags for the CODEX experiment

Carrier free antibodies were conjugated to selected barcodes using commercial reagents following Akoya Bioscience’s recommended protocols as described earlier ^93^. First, preservatives and other additives (like trehalose) were removed by using Amicon Ultra 30K centrifugation filters (Millipore, Cat. No. UFC503024). After multiple washes with PBS, the protein concentration was measured using an Implen nanophotometer with IgG mouse settings. 50 ug of protein was used for conjugation, diluted in 100 ul PBS.

For conjugation, Amicon Ultra 50K centrifugation filters (Millipore, Cat. No. UFC505024) (one for each conjugation) were washed with 500 ul filter blocking solution (Akoya Biosciences, Cat. No. 7000009 Part No 232113) and centrifuged at 12,000 x g for 2 min. The remaining solution was discarded and 50 ug antibody in a volume of 100 ul supplemented with PBS was added to the filter followed by centrifugation at 12,000 x g for 8 min. 260 ul antibody reduction master mix (Akoya Biosciences, Cat. No. 7000009 Part No. 232114 and Part No.232115) was added to the filter and incubated for 30 min. After centrifugation at 12,000 x g for 8 min, the filter was washed with 450 ul of conjugation buffer (Akoya Biosciences, Cat. No. 7000009 Part No. 232116). The barcode (one vial for each antibody) was resuspended in 10 ul nuclease free water (Ambion AM9938) and complemented with 210 ul of conjugation buffer. The mix was added to the filter and incubated for 2 hours. The filter was centrifuged at 12,000 x g for 8 min followed by washes (three times) with 450 ul purification solution (Akoya Biosciences, Cat. No. 7000009 Part No. 232117) and resuspended into 100 ul of antibody storage solution (Akoya Biosciences, Cat. No. 7000009 Part No. 232118). The antibody was collected by inverting the filter and centrifuging the content into a new tube at 3000 x g for 2 min. The collected conjugated antibody was stored at 4°C.

### GeoMX sequencing and analysis

The GeoMx DSP Spatial Proteogenomic Assay for both protein and RNA expression was performed following Bruker’s (formally NanoString) sample preparation protocol (MAN-10158). Briefly, SAVI patient and control lung slides were baked at 60°C for 40 minutes, followed by dewax and antigen retrieval performed on Leica BOND RX auto stainer using HIER2 (EDTA based antigen retrieval solution for 20 min at 100°C) and proteinase K (0.1 μg/ml 15minutes). RNA probes (NanoString GeoMx Whole Transcriptome Atlas Human RNA probes for NGS) were added for hybridization and incubated at 37°C for 18 h. The following day, slides were washed twice with a stringent wash (50% formamide in 4X SSC for 25 min each) followed by two washes with 2X SSC (2 min each). Slides were washed with 1X TBS for 5 minutes and then blocked with W buffer for 30 minutes. Subsequently, the antibody cocktail solution was prepared by mixing: GeoMX Human Protein Core for NGS, GeoMx Immune Cell Typing Panel Human Protein Module for NGS, anti-human CD68 (Clone SPM130) Alexa 488, anti-human CD45 Alexa 594, and anti-human aSMA Alexa 647 antibodies. Both, control lung and SAVI patient slides were incubated overnight in a 4°C refrigerator. The following day, samples were washed with TBS-T followed by fixation using 4% PFA and subsequently stained with SYTO13 as nuclear marker.

Slides were then washed and immediately loaded on the GeoMx Digital Spatial Profiler. Slides were imaged using a 20× objective, and ROIs from desired lung regions were selected based on CD68 and aSMA staining using the GeoMx software. After ROI selection, UV light was applied to each ROI to achieve subregion-specific indexing oligonucleotide cleavage. The released oligonucleotides were collected via a microcapillary and dispensed into 96-well plates. A total of 172 segments between the two samples were collected. Segments arise from 45 ROIs within control lung, and 84 ROIs in SAVI patient sample. All libraries were then prepared and sequenced on the Illumina Novaseq6000 per the manufacturer’s protocol performed by the NCI CCR Genomics Core. Fastqs for each ROI were then converted to Digital Count Conversion files using NanoString’s GeoMx NGS Pipeline v.2.3.3.10. All QC and downstream differential expression analyses were performed in NanoString DSP Data Analysis Suite.

### Generation of iPSC-derived endothelial cells (iECs) and isogenic controls

The SAVI patient-specific induced pluripotent stem cells (iPSCs) were generated from three patient fibroblasts and one patient’s PBMC using Cytotune 2.0 -iPSC Sendai reprogramming kit (ThermoFisher, Cat# A16517) that expresses Yamanaka’s transcription factors (Oct4, Sox2, Klf4, and C-Myc). Additionally, two isogenic control (iso-SAVI) iPSC lines were created from two patient-specific iPSCs (SAVI 1 and SAVI 2) with same *STING1* mutation (c.461, A>G) using CRISPR-Cas9 gene editing technology. Briefly, 0.8 million SAVI iPSCs were transfected with premixed 120pmol HiFiCas9 V3 (IDT #1081061), 200pmol STING-gRNA (Synthego), and 200pmol STING-ssODN (IDT) using Nucleofector 4D with buffer P3 and program CA-137 (Lonza #V4XP-3024) and then plated onto one well of rhLaminin-521 (Thermo Cat#A29249) coated 6-well plate with StemFlex (Thermo Cat#A3349401), 1x RevitaCell (Thermo #A2644501) and 1uM HDR enhancer V2 (IDT Cat#10007921). Transfected cells were cultured in 32°C incubator for 3 days before moving to 37°C incubator, and fresh medium was changed daily after transfection. Single-cell subcloning was done using single cell sorting to 1 cell/96-well Matrigel coated plate with StemFlex and CloneR2 (Stem Cell Technologies, Cat#100-0691). 10 days later, single-cell clones were picked and confirmed by genomic PCR using STINGNHEJ-F2/R1 primers, Sanger sequencing, and ICE analysis (Synthego). iPSC lines from three healthy volunteers (HC) were also generated. These iPSC lines were differentiated into endothelial cells (iECs) following our previously published protocol ^94^.

In brief, iPSCs from HC, SAVI, and iso-SAVI groups were seeded on Matrigel (BD Biosciences, Cat#354230)-coated plates at low density and supplemented with TeSR-E8 (Gibco cat. no. A1517001). After 24 hours, the cells were induced toward mesoderm progenitors using mesoderm differentiation medium for six days. Subsequently, CD31 positive cells from the induction culture were enriched using CD31 magnetic beads (Miltenyi Biotec Cat#130091935), and then further seeded at approximately 2000 cells/cm2 on Corning® BioCoat® Collagen I plates (Corning, Cat#356450) and cultured in a 1:1 mixture of EGM2 growth media (EGM^TM^-2, Lonza, Cat#CC-3162) and Human Endothelial-SFM (Cat#11111044 Fisher Scientific). The media was changed every other day. Once reaching 80-90% confluence, iECs at passage 1 were used for further analysis. Meanwhile, cells were passaged using TrypLE™ Express Enzyme (Invitrogen Cat#12605010) and expanded under the same culture conditions until passage 5.

### Flow Cytometry Analysis of iECs

iECs derived from HC, SAVI and iso-SAVI groups at different passages or different treatment groups were collected for flow cytometry analysis of endothelial surface marker CD144 (VE-cadherin) and CD31. Cells were dissociated with TrypLE™ Express Enzyme, resuspended in flow cytometry analysis buffer (PBS with 0.5% BSA) and further stained with PE-conjugated CD144 (BD Cat#560410) and APC-conjugated CD31 (Biolegend Cat# 303116) for 30 min on ice following the manufactory instruction. After wash with PBS, cells were analyzed by MACSQuant Flow Cytometer (Miltenyi Biotec, Bergisch Gladbach, Germany and the results were analyzed with MACSQuant analysis software. The percentages of positive cells are determined by comparison with isotype controls that establish the background levels due to nonspecific staining.

### Tube formation assay

The tube formation assay was performed as previously described ^94^. Briefly, 3000 iECs per well were plated on a 96-well plate coated with 65μl of Matrigel Basement Membrane Matrix (BD Biosciences, Cat#354234) and cultured for 6 hours. After removing the medium from the plate, the cells were labeled by adding 100μl per well of 8μl/ml BD calcein AM (BD Biosciences, Cat#564061). Following a 30-40 minute incubation, the cells were photographed using a fluorescent microscopy system. The number of branch points and tubule lengths were quantified using ImageJ.

### Low-density lipoprotein (LDL) uptake assay

A low-density lipoprotein (LDL) uptake assay was performed as described previously ^94^. Briefly, the iECs were incubated with 10 μg/ml acetylated LDL (Ac-LDL) labeled with 1,1′-dioctadecyl-3,3,3′,3′-tetramethylindo-carbocyanine perchlorate (DiI-Ac-LDL) (Invitrogen, Cat#L3484) for 4 h at 37 °C. The DiI-Ac-LDL uptake was assessed and quantified by fluorescent microscopy. Subsequently, cells were dissociated into single cells with TrypLE TM Express Enzyme. The data acquisition was performed on a MACSQuant Flow Cytometer (Miltenyi Biotec, Bergisch Gladbach, Germany) and the results were analyzed with MACSQuant analysis software.

### Cell immunofluorescence staining

Cells were fixed with 4% paraformaldehyde for 10 minutes and washing with PBS, blocking solution (5% normal donkey serum in PBS with 0.3% Triton-X100) was applied for 1 h. at RT. The primary antibody VE-Cadherin (1:100 Santa Cruz Cat# sc-9989), CD31 (1:50, DAKO Cat#M0823), IRF3 (1:200, CST,Cat#11904S), STAT3 (1:800, CST Cat#9139S) was applied overnight at 4°C, followed by the corresponding secondary fluorescence-conjugated antibody 1:400 (Invitrogen) for one hour at RT. DAPI (1:1000, Invitrogen) was used for cell nuclei detection. Cells were imaged using a Zeiss inverted fluorescence microscope (Zeiss Axio Vert A1, Germany). These antibodies are listed in the Key Resource table.

### Human Primary Lung Microvascular Endothelial Cells (HLMEC) culture and stimulation

HLMEC cells (Lonza) were cultured in EGM2^MV^ growth media (Lonza). Cells from passages 3-6 were utilized for experiments. The cells were plated at a density of 60,000 cells per well in a 12-well plate for immunofluorescence staining, and at 100,000 cells per well in a 6-well plate for protein or RNA isolation. HLMEC cells were treated continuously with 20 μg/ml of the STING ligand Cyclic [G(2’,5’)pA(3’,5’)p] (2’3’cGAMP) (Invivogen, Cat# tlrl-nacga23-5) and/or 1000 units/ml IFNβ (human IFN beta 1a, PBL Assay Science Cat# 11410-2) for 8 hours, 1, 3, and 5 days. By different time point, cells were either fixed for immunofluorescence staining or harvested for subsequent analysis by Western Blot, RT-PCR, RNA-seq, or ATAC-seq.

### siRNA transfection in HLMEC

siRNA plasmids for control (sc-37007), STAT3 (sc-29493), and STAT1 (sc-44123) were purchased from Santa Cruz. HMVEC-L cells were plated at a density of 150,000 cells per well in a 6-well plate. The following day, cells were transfected with siRNA plasmids using Lipofectamine RNAiMAX transfection reagent (Thermo Fisher Scientific, Cat# 13778075) following the manufacturer’s instructions. Knockdown efficiency was evaluated 48 hours post-transfection. Subsequently, HLMECs were treated with 20 μg/ml of the STING ligand Cyclic [G(2’,5’)pA(3’,5’)p] (2’3’cGAMP) (Invivogen, Cat# tlrl-nacga23-5) and/or 1000 units/ml IFNβ (human IFN beta 1a, PBL Assay Science, Cat# 11410-2). At different timepoint (4 hours or 3 days) after stimulation, cells protein were harvested for Western Blot analysis.

### shRNA knockdown or cDNA overexpress in SAVI iECs

shRNA (h) lentiviral particles for STAT3 (Cat#sc-29493-V), STAT1 (Cat#sc-44123-V), STING1 (previously TMEM173) (Cat#sc-92042-V), and scrambled controls (sc-108080) were purchased from Santa Cruz. Lentiviral particles for SOX18 (NM_018419) Human Tagged ORF Clone and Lenti ORF control were purchased from OriGene (Cat#SKU RC210323L4V, SKU PS100093V). Passage 1 SAVI iECs were seeded in a 6-well plate at a density of 150,000 cells/well. When the cells reached 70% confluence (the next day), shRNA or cDNA lentiviral particles, including 100,000 infectious units of virus (IFU), were applied to the cells with Polybrene (sc-134220) at a final concentration of 5 µg/mL in 1 mL of iEC culture medium overnight. An additional 1 mL of fresh iEC culture medium without Polybrene was added to the well for another day. STAT3, STAT1, and STING1 shRNA knockdown efficiency or SOX18 cDNA overexpression was assessed 48 hours after infection. Due to the extended duration of the experiment, 1 µg/mL puromycin was maintained in the culture to select only the infected cells until the endpoint of the experiments, typically at passage 5. The iECs were passaged and expanded as previously described.

### Western Blot

Western blot samples were lysed in CHAPS buffer containing protease and phosphatase inhibitor cocktails (Invitrogen). For each sample, 20 μg of total protein was analyzed by Western blot using a Bio-Rad 4–20% Mini-PROTEAN TGX Gel and subsequently transferred onto nitrocellulose membranes. After blocking with LI-COR blocking buffer, the membranes were incubated with primary antibodies for VE-cadherin (1:1000, CST, Cat#2500S), CD31(1:1000, DAKO, Cat#M0823), SMA (1:2000, Abcam, Cat#ab7817), SM22 (1:1000, Abcam, Cat#ab14106), SOX18 (1:100, Santa Cruz, Cat#sc-166025), SLUG (1:1000, CST, Cat#9585S), P-STAT3^Y705^ (1:1000, CST, Cat#9145S), STAT3 (1:1000, CST, Cat#9139S), P-STAT1^Y701^ (1:1000, CST, Cat#9167S), STAT1(1:1000, CST, Cat#9172S), P-STING^S366^ (1:1000, CST, Cat#19781S), STING (1:1000, CST, Cat#13647S), P-TBK1^S172^ (1:1000, CST, Cat#5483S), TBK1 (1:1000, CST, Cat#3504S), P-IRF3^S396^ (1:1000, CST, Cat#4947S), and IRF3 (1:1000, CST, Cat#11904S) overnight at 4°C; these antibodies are listed in the Key Resource table. A secondary fluorescence antibody (1:10,000 LI-COR) was applied for 1 hour at room temperature. After washing three times with PBST, the signal was detected and quantified by the LI-COR Odyssey CLx Imager and Image Studio^TM^ software.

### RNA extraction, reverse transcription and real-time PCR

RNA was isolated from cultured cells using the RNeasy kit (Qiagen, Cat#74106) according to the manufacturer’s instructions. RNA was quantified using a Nanodrop NP-1000 spectrometer (Wilmington, DE). Reverse transcription was performed using the SuperScript™ III Reverse Transcriptase kit with dNTP, Random Hexamer (Invitrogen, Cat#18080093), and RNAase inhibitor (New England Biolabs) . Reaction conditions were: 5°C for 10 min, 48°C for 30 min, 95°C for 5 min and 15°C forever. One μg RNA was used per reaction. Subsequently, real-time PCR was performed using the iQ Syber Green kit (Bio-Rad, Cat#1708880) according to the manufacturer’s protocol. 18S RNA served as control. Quantitative PCR conditions were as follows: 95°C for 5 min, with 39 cycles consisting of 95°C for 45 sec, 57°C for 45 sec, 72°C for 45 sec, read at 78°C, and 95°C for 45 sec. Melting curve determination ranged from 72°C to 95°C with 0.2°C increments. RNA expression was analyzed using the ΔΔCt method. qPCR primer sequences are listed in Key Resource table.

Specific for Figure 1G: Two(2) control and Six(6) SAVI patients fibroblast were cultured and stimulated with 20ug/ml cGAMP (Invivogen, Cat. No.-tlrl-nacga23) and 10ng/ml TGFβ (R&D Systems, Cat. No.-240-B-010/CF) up to 72hours. Total RNA was isolated by RNeasy kit (Qiagen, cat no-74106) and cDNA were prepared using SuperScript IV First-Strand Synthesis kit (Invitrogen, Cat.No.-18091050) according to the manufacturer’s instructions. mRNA expression of target genes (Mentioned in the table below) were performed by real time PCR using TaqMan gene expression Master Mix (Applied Biosystem, Cat. No-4369016.). HPRT1was used as a control. The target genes were amplified using TaqMan PCR primers showed in Key Resource table.

### Bulk RNA sequencing

Bulk RNA-seq analysis was conducted on samples of iPSC-derived endothelial cells (iECs) from healthy controls (HC), SAVI patients, and isogenic controls for SAVI (iso-SAVI),. RNA was extracted using a combination of TRIzol and RNeasy mini kit (Qiagen). The integrity of the total RNA was evaluated using automated capillary electrophoresis on a Fragment Analyzer (Roche). For all samples with an RNA quality indicator (RQI) greater than 8.0, 500 ng of total RNA was used for library preparation using the TruSeq Stranded mRNA Library Preparation Kit (Illumina). The sequencing libraries were quantified by PCR using the KAPA Library Quantification Kit for NGS (Kapa) and evaluated for size distribution and absence of adapter dimers using a Fragment Analyzer. The libraries were then pooled and clustered on a cBot2 (Illumina) with a HiSeq 3k/4k PE Cluster Kit, followed by sequencing on a HiSeq 3000 platform (Illumina) under conditions for 75 bp paired-end reads and a 7 bp indexing read.

### Bulk RNA-Sequencing analyses

Bulk RNA-seq samples were processed using the RNA-seek pipeline (https://zenodo.org/doi/10.5281/zenodo.4901427). Initially, adapter sequences were removed and quality trimming was performed using Cutadapt v1.18 (http://dx.doi.org/10.14806/ej.17.1.200). The processed reads were then aligned to the human GENCODE v43 primary assembly (GRCh38) using STAR v2.7.6a (https://doi.org/10.1093/bioinformatics/bts635) in two-pass basic mode. Gene and transcript quantification were subsequently conducted using RSEM v1.3.0 (https://doi.org/10.1186/1471-2105-12-323), based on the GENCODE v43 ‘PRI’ gene annotations. Differential expression analyses were performed using DESeq2 v1.42.1 (https://doi.org/10.1186/s13059-014-0550-8) within the R v4.3.3 statistical software environment (https://doi.org/10.59350/t79xt-tf203). Integer-based gene counts were derived from the RSEM gene count estimates using tximport v1.30.0 (https://doi.org/10.12688/f1000research.7563.1). Genes were considered differentially expressed if they had a false discovery rate-adjusted p-value of less than 0.05 and an absolute fold-change greater than 1. Additionally, genes with no expression (fewer than 1 read) in over half of the samples in each comparison were disregarded.

Gene Ontology (GO) enrichment tests were conducted on differentially expressed gene lists using clusterProfiler v4.10.1 (https://doi.org/10.1016/j.xinn.2021.100141), focusing on biological process ontology terms. GO terms with a Benjamini-Hochberg adjusted p-value of less than 0.01 were considered significantly enriched.

Ingenuity Pathway Analyses (IPA) were conducted using the IPA software tool (https://doi.org/10.1093/bioinformatics/btt703). As part of the core analyses, IPA upstream analyses were performed on differentially expressed genes, incorporating FDR-adjusted p-values and log2-normalized fold-changes. Upstream regulators were considered activated if they had a Z-score greater than 2 and a p-value of differential gene overlap less than 0.0-1, while inhibited upstream regulators were identified by Z-scores less than -2.

### ATAC-seq (Assay for Transposase-Accessible Chromatin with high-throughput sequencing) library preparation and sequencing

ATAC-seq sample preparation was followed and modified from Chang lab-Omni-ATAC protocol^95^. Frozen cells (iECs from HC and SAVI at P1 and P3, HLMEC 8 hours after stimulation with or without cGAMP or IFNβ) were gently thawed, counted and resuspended in PBS. 50000 cells per sample were lysed with ATAC-seq Lysis Buffer (50 ul cold ATAC-Resuspension Buffer (RSB) (10 mM Tris-HCl PH 7.4, 10 mM NaCl, 3 mM MgCl_2_ in sterile water) containing 0.1% NP40, 0.1% Tween20, and 0.01% Digitonin) for 5 min and washed with RSB buffer with 0.1% Tween-20. Resuspend cell pellet in 100 ul of transposition mixture by pipetting up and down 6-12 times.

Transposition 50 ul mix = (25 ul 2x TD buffer (20 mM Tris-HCl PH 7.4. 10 mM MgCl_2_, 20% Dimethyl Formamide in sterile water), 2.5 ul transposase (100nM final), 16.5 ul PBS, 0.5 ul 1% digitonin, 0.5 ul 10% Tween-20, 5 ul H2O). Incubate reaction at 37°C for 30 minutes in a thermomixer with 1000 RPM mixing. Clean up reaction with a Qiagen MinElute PCR Purification Kit (Qiagen Cat# 28004). Set up the PCR reactions by adding unique i5 and/or i7 index primer. Pre-amplification of transposed fragments was used to determine the required number of additional cycles to amplify ^96^.After final amplification and cleanup, DNA fragmentation was determined by Bioanalyzer. DNA was eluted in 20 ul volume at -20 degree for further sequencing. The library concentration was quantified by using the KAPA Library Quantification Kit (cat# KK4854).

Sequenced libraries were trimmed for adapters using the trim galore wrapper of the Cutadapt and FastQC packages (version 0.6.10), and reads were mapped to the hg38 genome using the Burrows-Wheeler aligner (version 0.7.17) ^97,98^. Reads were filtered for a map quality of greater than 40 using the samtools package (version 1.11) and PCR duplicates were removed using the Mark Duplicates utility within Picard tools (version 2.27.4) ^99^. Peaks were called using the Genrich package with a p-value threshold of 0.005 and a q-value threshold of 0.01. Differential accessibility was called between the different experimental conditions using the DiffBind package (version 3.1.4) with a threshold of FDR < 0.01 and a |fold change| > 0.5 ^100^. Transcription factor motif enrichment was performed in the differentially accessible regions using the Monalisa (version 1.10.1) package and the motifs found within the JASPAR 2024 database ^101,102^. Genes were annotated to the differentially accessible regions using the default settings of the GREAT web browser tool (version 4.0.4) ^103^. Gene ontology pathway enrichment metrics were also obtained from the GREAT web browser tool. Permutation tests for enrichment of overlap between ATAC-seq peaks and HUVEC ChIP-seq peaks were performed using the RegioneR package (version 3.22)^104^ with 500 random permutations. ChIP-seq datasets used for the permutation tests include the HUVEC SOX18 ChIP-seq from Overman et al 2017, HUVEC ERG ChIP-seq from Kalna et al 2019, HUVEC BRD4 ChIP-seq from Brown et al 2014^54^, and the HUVEC ENCODE datasets (JUN: ENCFF624TOT, FOS: ENCFF972ZIV, GATA2: ENCFF987YIJ) ^57,105–107^. For the tests to compare the enrichment of the TNF-stimulated BRD4 peaks and the unstimulated BRD4 peaks from Brown et al 2014 study, a greater number of permutations was used (1000), and a differential permutation test was also implemented. This differential permutation test assessed the degree to which the difference in overlaps observed between the different BRD4 peak sets and the reference region set was greater than a randomly generated null distribution that represents the expected number of differences in the number of overlaps between the TNF-stimulated BRD4 peaks and the reference set compared to the number of overlaps between the unstimulated BRD4 peaks and the reference set. A significant result in this test would mean that the TNF-stimulated BRD4 peaks intersect the reference region set more times than the unstimulated peaks to a degree greater than would be expected by random chance based on the size and number of regions in all of the region sets.

## Supporting information

Supplemental Fig S1-S6 with figure legends and Supplemental table S1

## Materials availability

This study did not generate new unique reagents.

STING inhibitor IFM35883 Provided by IFM Therapeutics (IFM) under a government CRADA.

## Data and code availability

Raw data of RNA-seq and ATAC-seq experiments derived from SAVI patients were deposited with dbGAP number phs001946.v3.p1. Healthy control data were generated from specimens collected prior to implementation of the 2015 NIH Genomic Data Sharing Policy and therefore cannot be deposited in dbGaP. Aggregate statistics and processed results are provided in the Supplementary Materials, and requests for individual-level control data may be directed to [corresponding author email] for case-by-case review in accordance with IRB and consent requirements.

All data that support the findings of this study are available from the Lead contact upon reasonable request.

This paper does not report original code.

Any additional information required to reanalyze the data reported in this paper is available for the lead contact upon request.

## Lead contact

Further information and requests for resources and reagents should be directed to and will be fulfilled by the lead contact, Dr. Raphaela Goldbach-Mansky.

## Key resources table

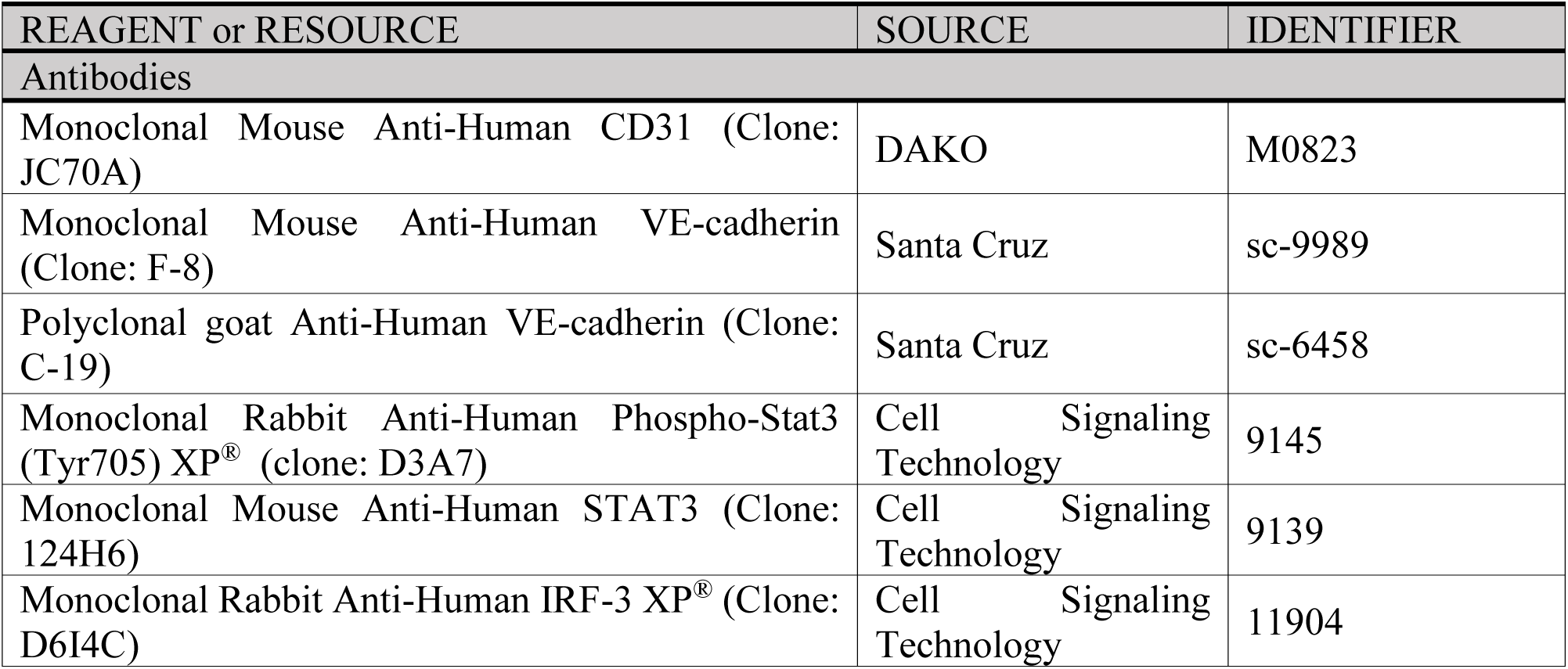
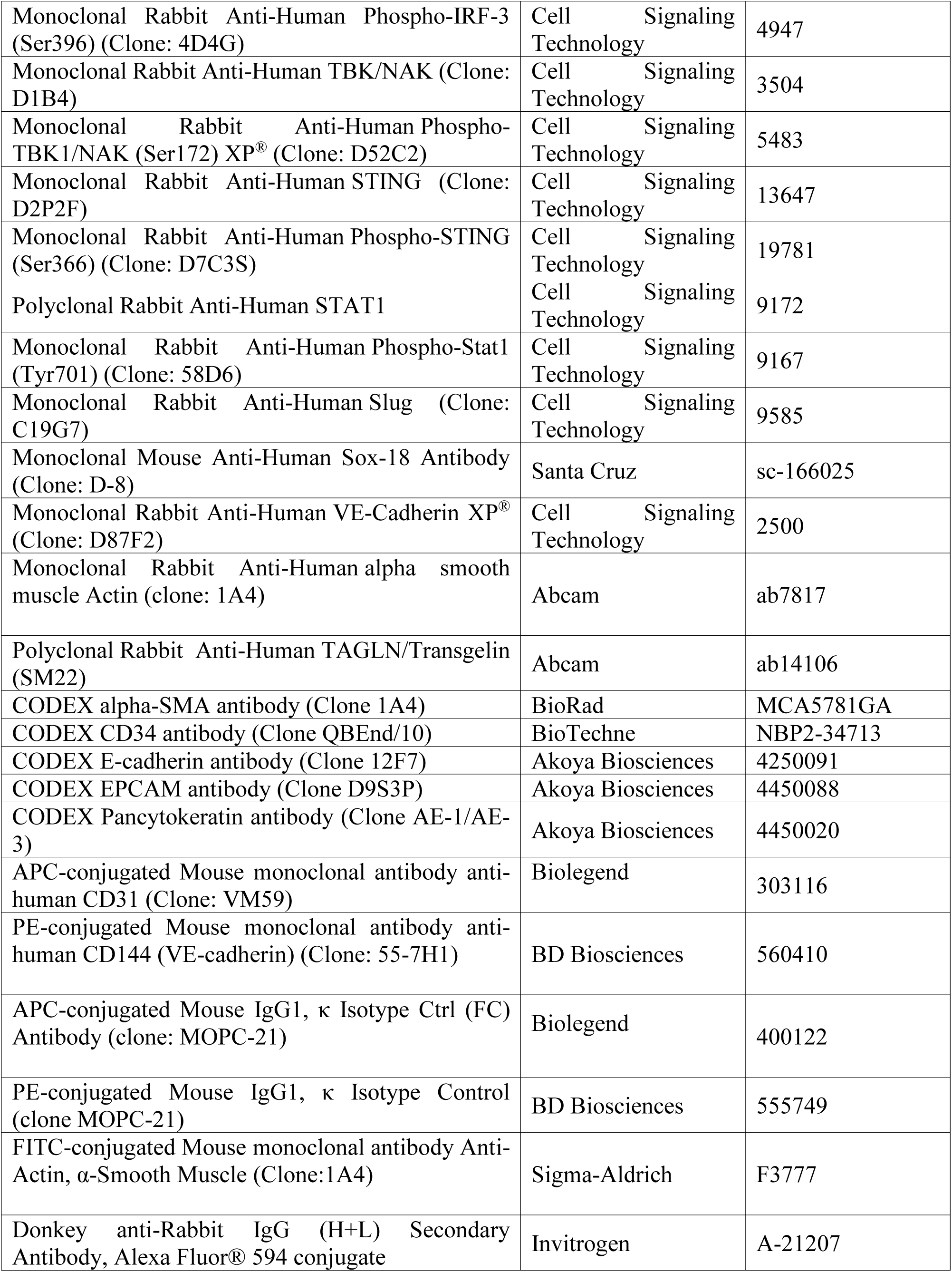
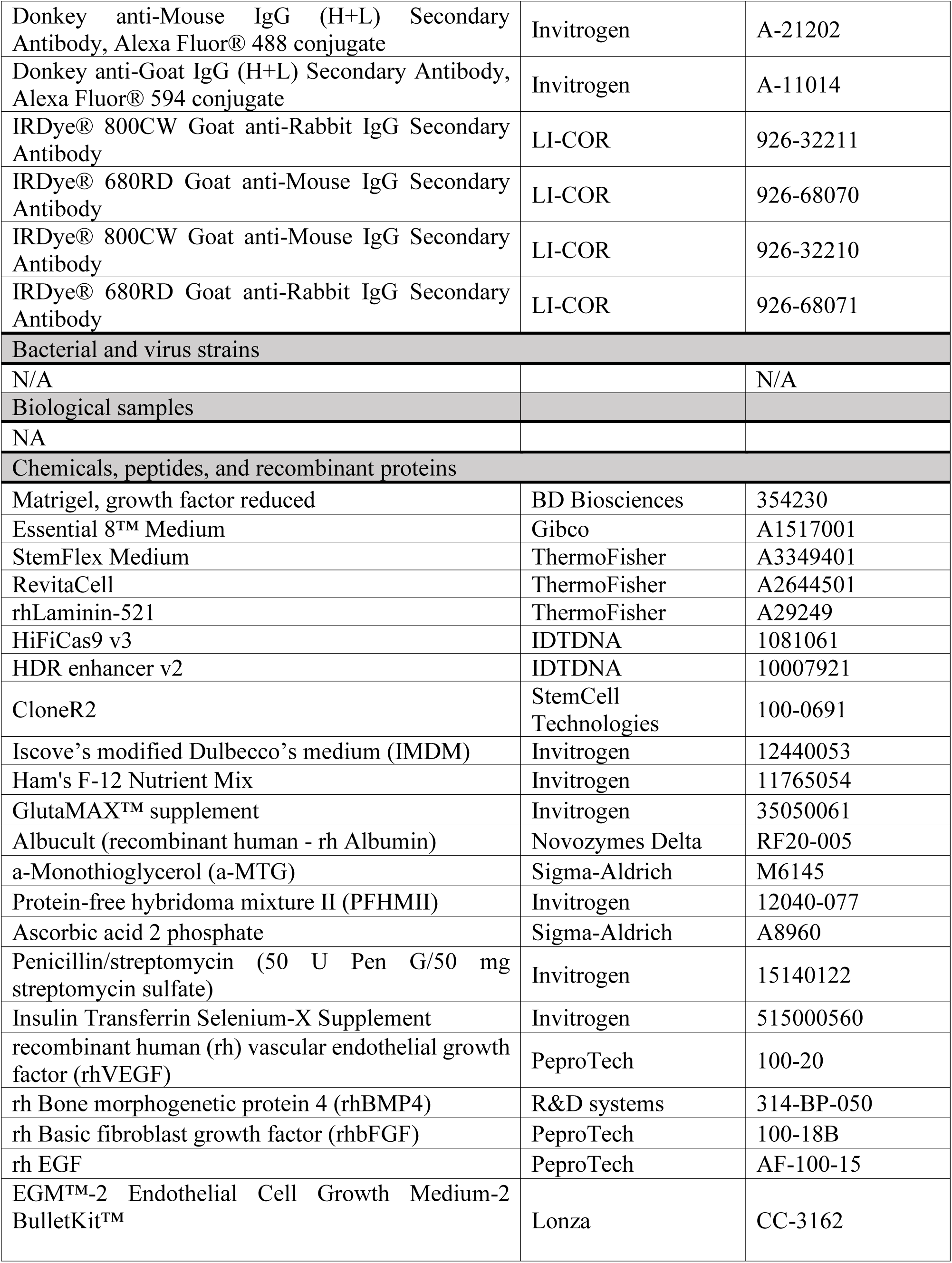
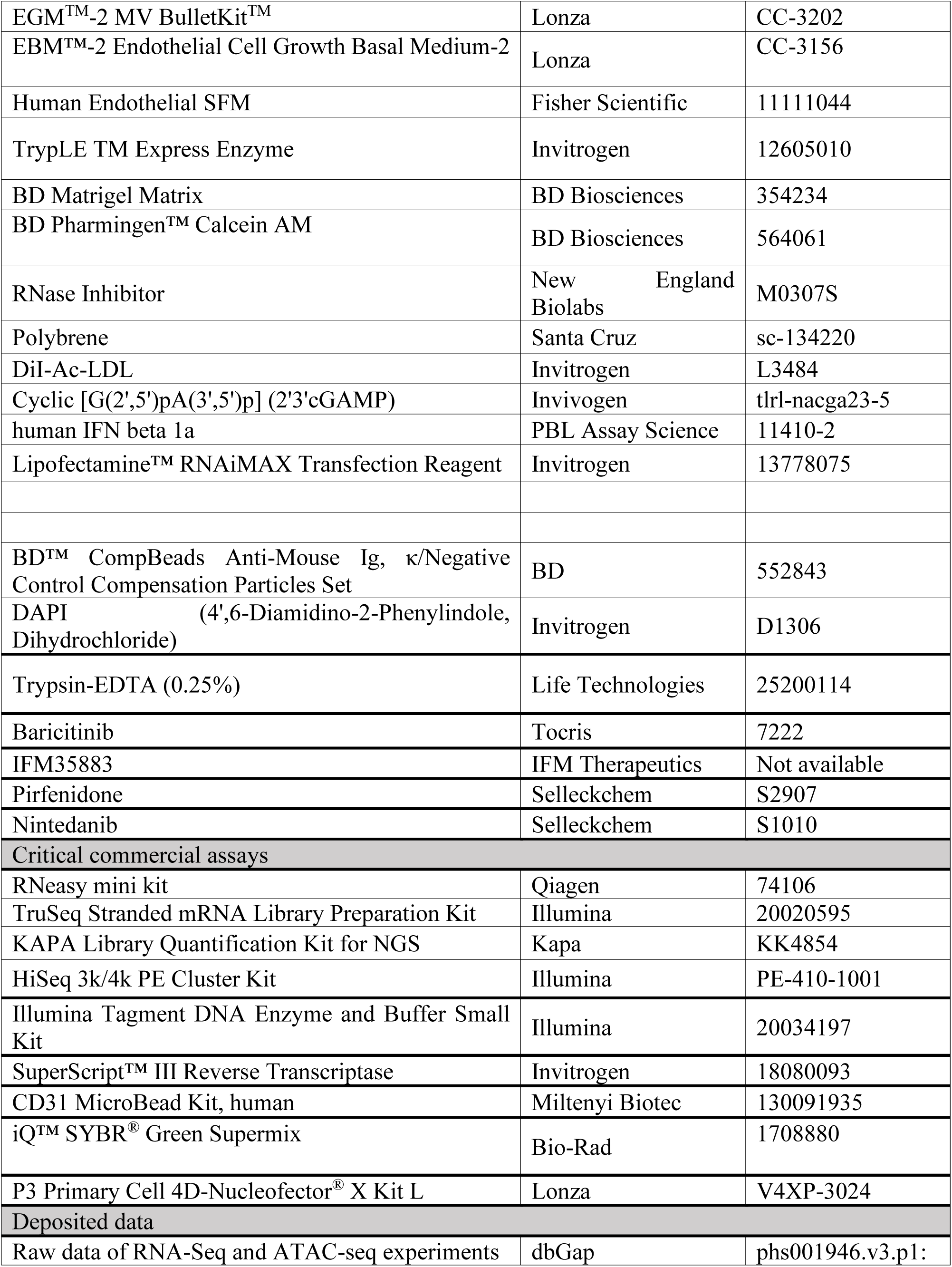
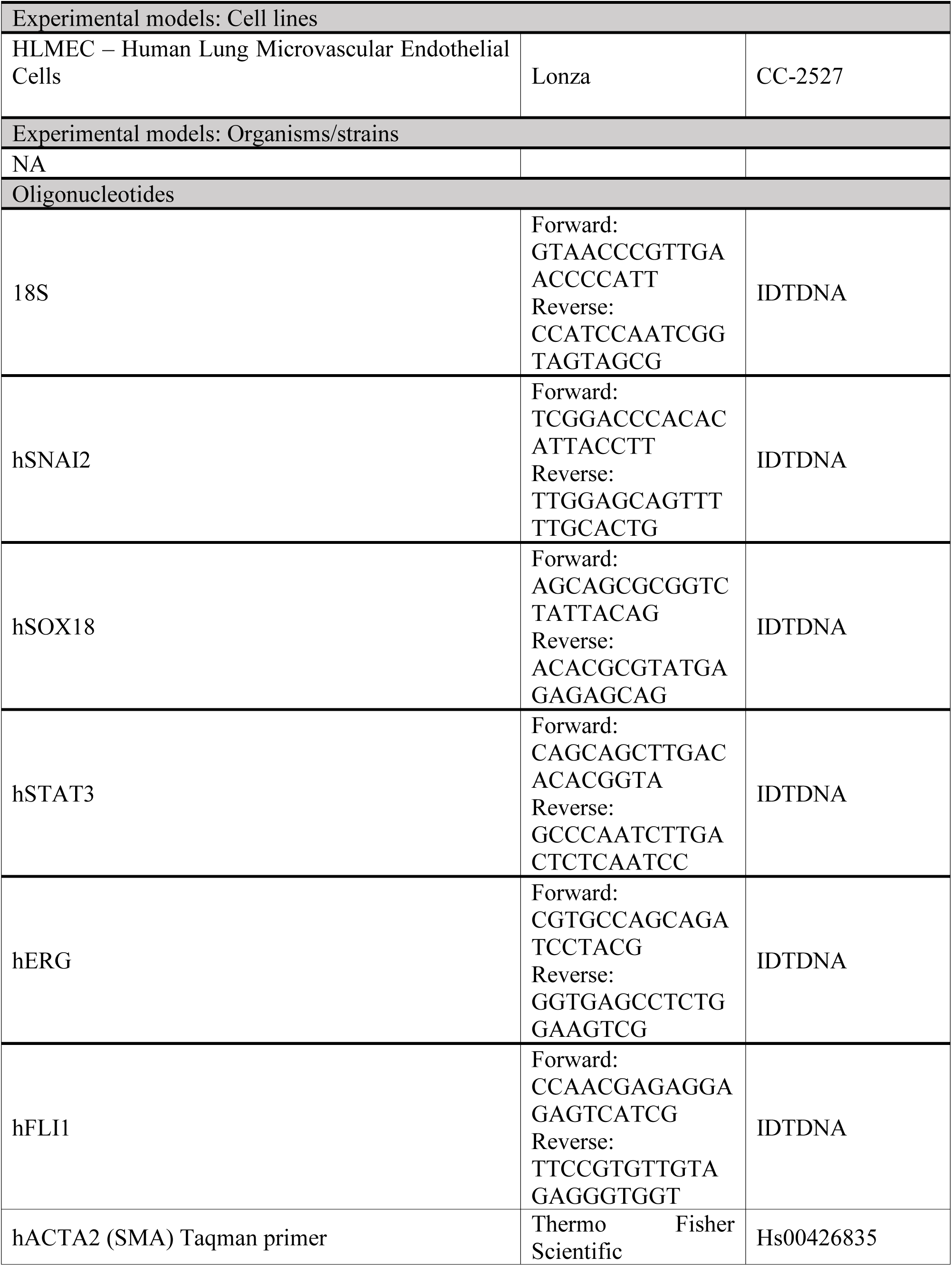
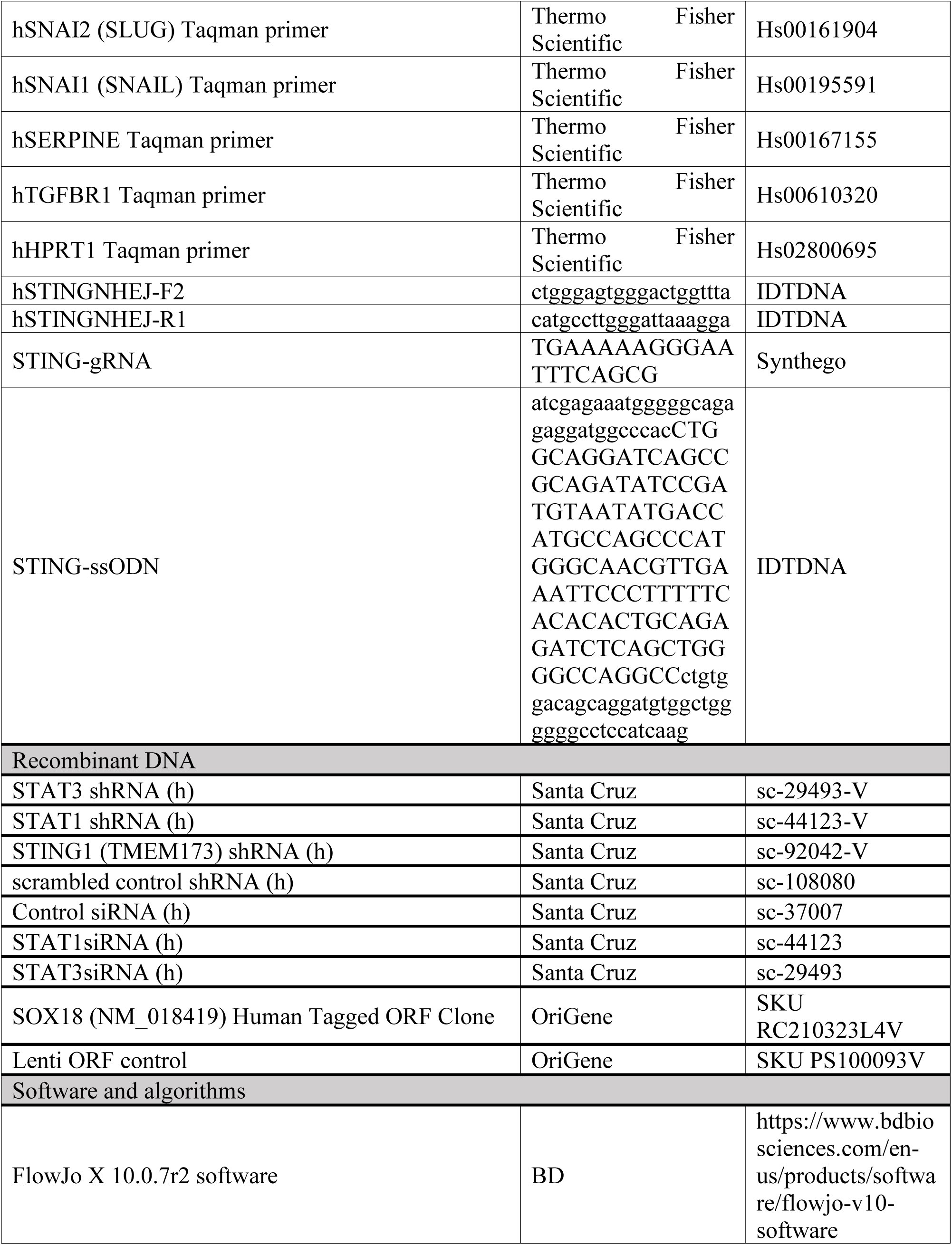
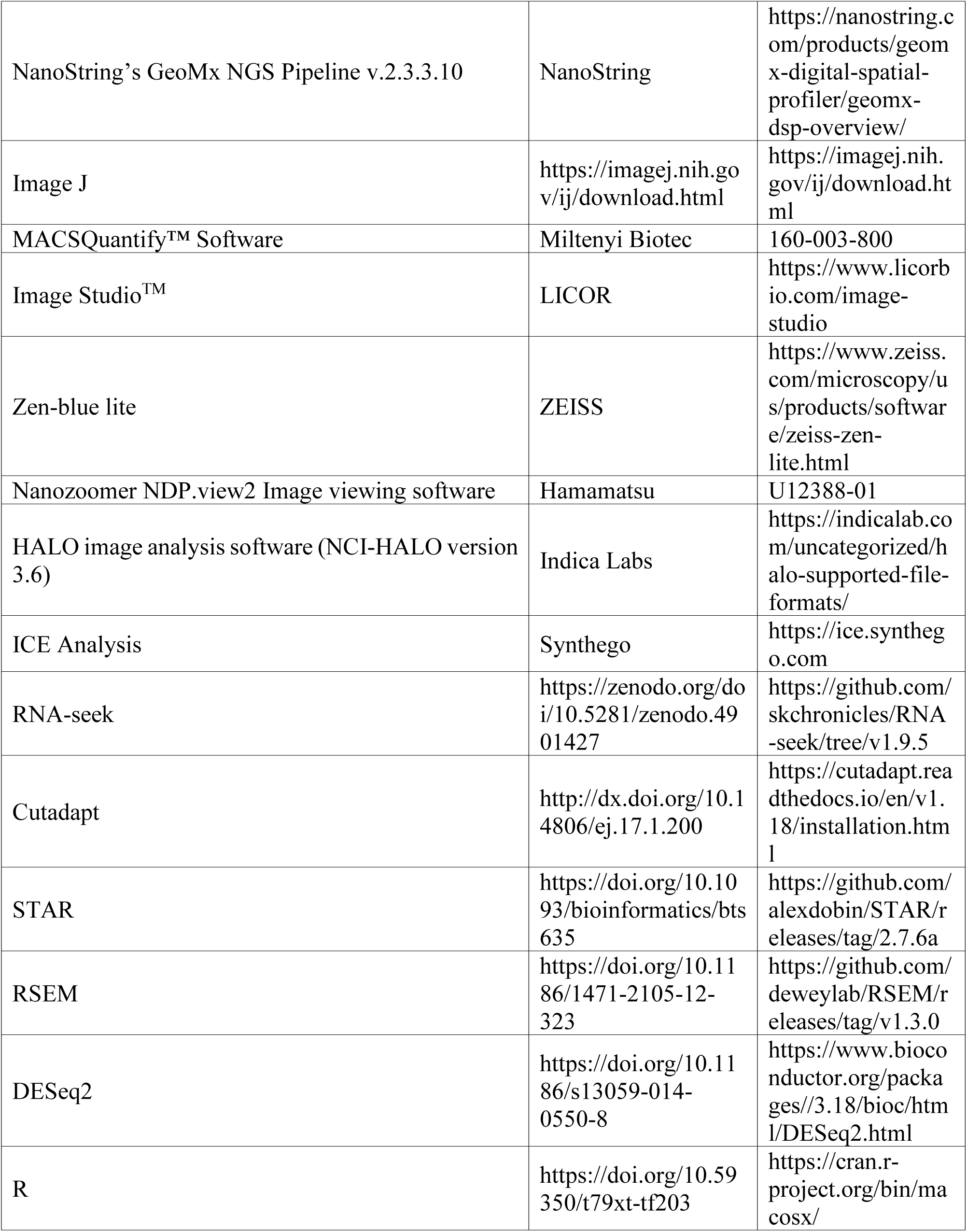
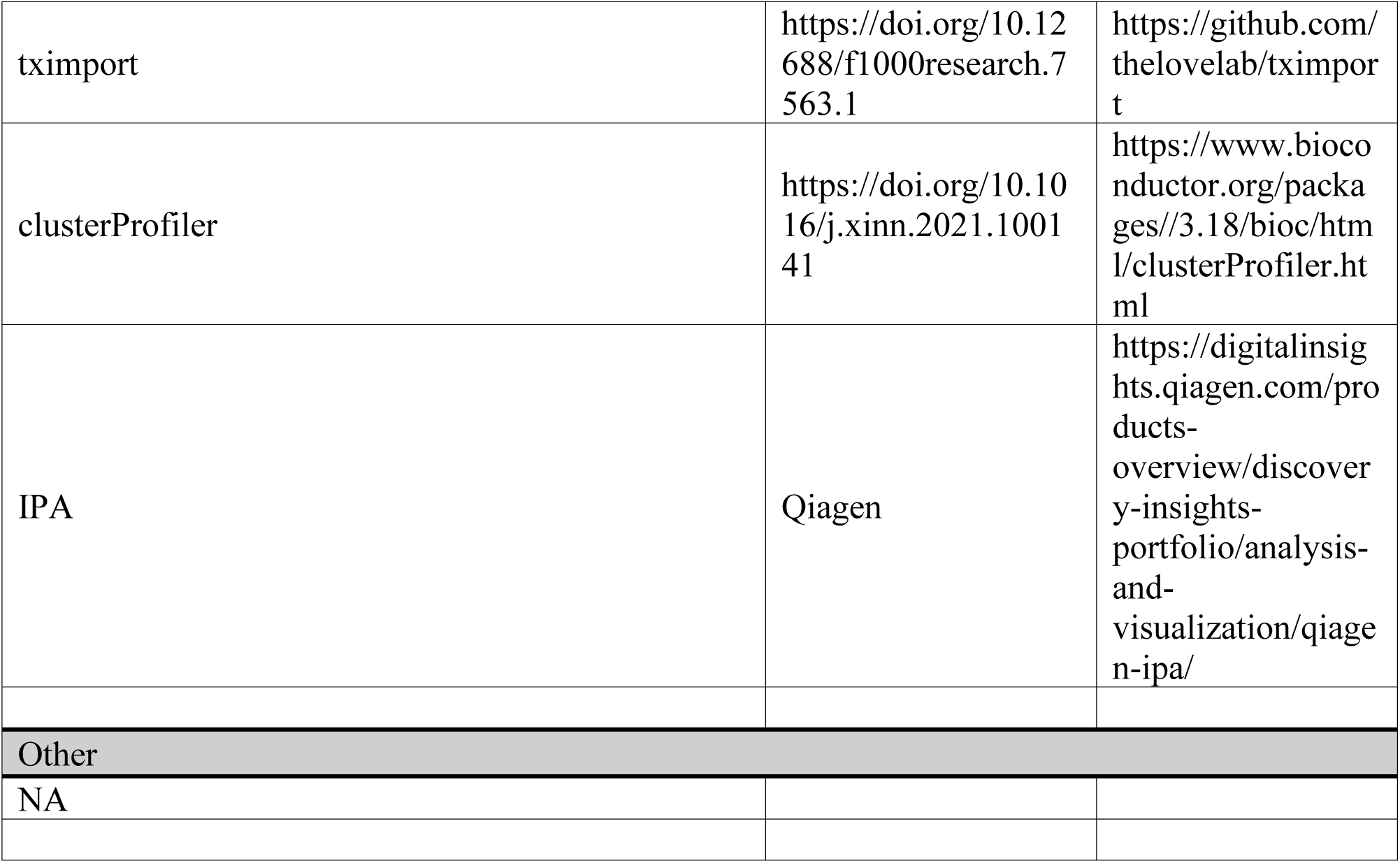

## Acknowledgments

This research was supported by the Intramural Research Program (IRP) of the National Institute of Allergy and Infectious Diseases (NIAID) and National Heart, Lung, and Blood Institute (NHLBI), National Institute of Health (NIH). The contributions of the NIH authors are considered Works of the United States Government. The findings and conclusions presented in this paper are those of the authors and do not necessarily reflect the views of the NIH of the U.S. Department of Health and Human Services. We would like to thank Drs. Maria Romero (CV Path Institute), Dr. Renu Virmani (CV Path Institute), and Dr. Zu-Xi Yu (NHLBI Pathology Core) for their help with reviewing the lung biopsies and for helpful discussions; Dr. Stefan Muljo (NIAID) for the early assistance with advice and epigenomic sequencing and Andrew Oler (NIAID Bioinformatics and Computational Biosciences Branch (BCBB)) for his assistance with preliminary RNA seq data analysis in HLMECs; Dr. Elizabeth A. Conner (NCI CCR core) for her help with CODEX and GeoMX experiments; Dagmar Bacikova for helping with RNA sequencing in the American Genome Center, Uniformed Services University of the Health Sciences; and Dr. Elisa Ferrante and Dr. Cornelia Cudrici (NHLBI) for clinical protocol consultant; Louise Malle for early preliminary experiments, and Dr. Bradley Peterson, Drs. Megan Curan and Donna Curtis for excellent patient care.

## Author contributions

D.Y: experiments design and conduction, data interpretation, data analysis, figure design, manuscript writing, manuscript revision; G.C: generation of iPSCs, iEC differentiation and characterization, data analysis. figure design; S.G: fibroblast experiments; A.dJ: histology scoring, sample coordination; A.K.M: clinical data interpretation, conducting experiments and manuscript writing; C.McN: Statistical analysis bioinformatics support; A.M: Bioinformatics support; J.W: HLMEC experiments; N.K: CODEX staining and spatial transcriptomics; M.H: spatial transcriptomics; J.Z: design and generation of SAVI isogeneic iPSC line; K.L: culture and characterization of SAVI isogeneic iPSC line; G.S.: RNA and ATAC sequencing; C-C.L: lung histology scoring; Y.Z.: experiments design and conduction, bioinformatics support; S.A: Patient data collection; L.F: imaging analysis; Q.Y: iPSC maintenance and differentiation; Bin Lin : early experiments; B.L: patient data and care, manuscript editing; B.B: patient data and care; A.R: patient data and car;. G.S: patient data and care; D.R.L: patient data and care; B.P: pathology sample acquisition; S.O: patient data and care; A.B: patient data and care; M.W.: GeoMX sequencing technique support; D.T.: GeoMX sequencing bioinformatic support; N.D: bioinformatics support; H.M: histology support and IPF sample contribution; J.C.K: manuscript editing and discussion of experimental details; C.D: sequencing and data analysis support; M.B: Study design, figure design, data interpretation, manuscript writing and revision, funding acquisition; R.G-M: Study design, figure design, data interpretation, primary manuscript writing and revision. All authors improved and accepted the final version of the manuscript.

## Declaration of interests

RGM received the STING inhibitor compound used in the studies under a government approved CRADA with IFM. All other investigators declare no conflict of interest.

## Supplemental information

Document S1. Figures S1–S6 and Table S1

## Notes

### Competing Interest Statement

The authors have declared no competing interest.

