## Supplemental Fig S1-S6 with figure legends and Supplemental table S1 for "STING–STAT3–SOX18 Axis Drives EndMT and Epigenetic Reprogramming in SAVI Lung Fibrosis"

**Figure S1**

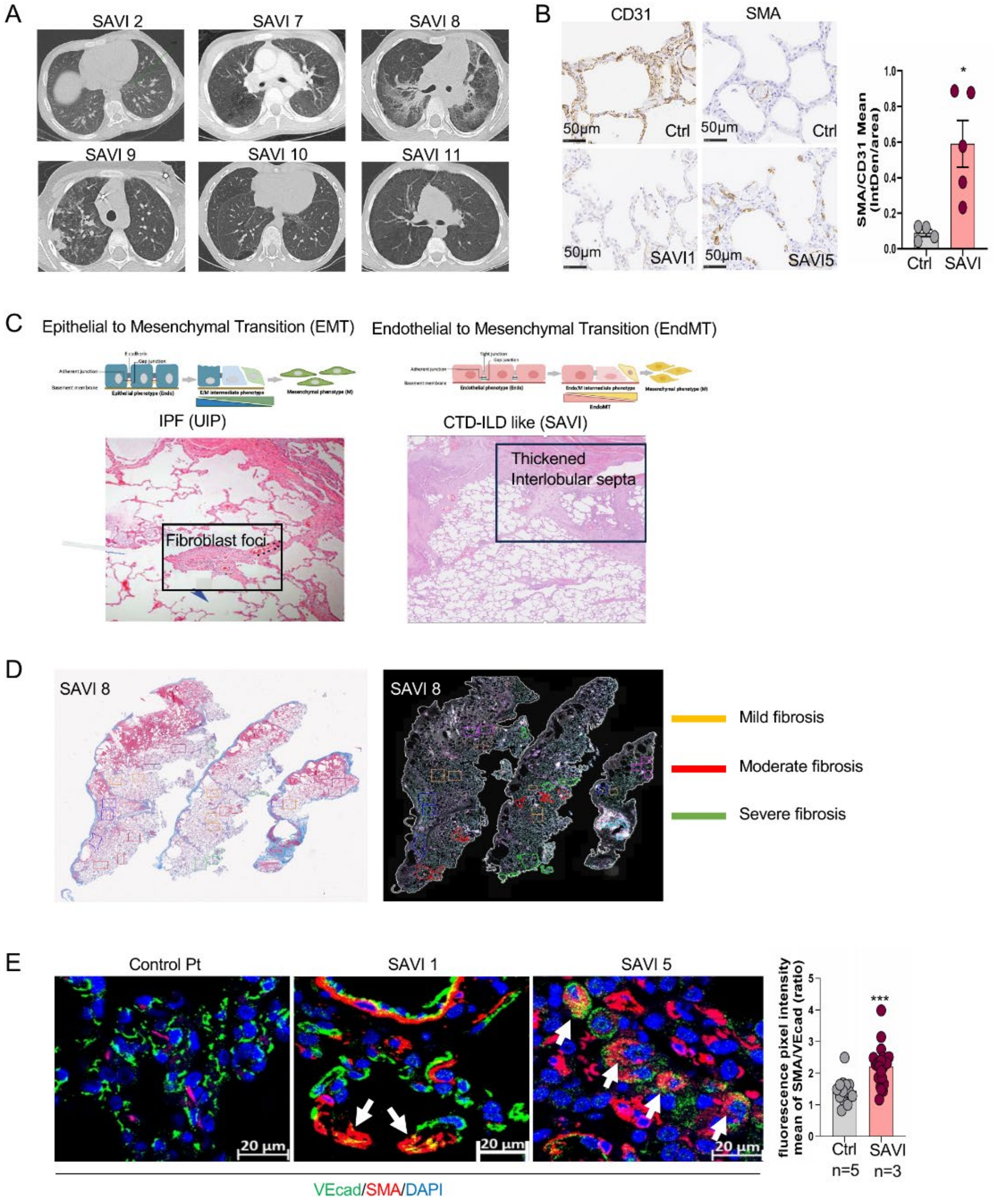


**Figure S1: Histopathological features of pulmonary fibrosis associated with SAVI reveal microvascular damage and fibrosis, Related to Figure 1.**

(A) CT Chest images from SAVI patients exhibit a variety of interstitial lung disease (ILD). These images were taken from 6 SAVI patients (SAVI 2, 7-11). Some SAVI patients do not exhibit any ILD findings on CT (not featured) while others exhibit findings such as interspersed cystic lesions, heterozygous peripheral ground glass opacities and consolidations, mosaic attenuation reflecting air trapping/small airways disease, and reticular opacities including interlobular and intralobular septal thickening. As the ILD becomes more advanced, architectural distortion will develop, including fibrotic reticulations, traction bronchiectasis, volume loss, and eventually honeycombing. Supplemental to Figure 1A.

(B) Immunohistochemistry staining for the endothelial marker CD31 and the mesenchymal marker SMA in the lung alveolar area from control and SAVI patients. Positive cells are stained brown, and nuclei are counterstained blue with Hematoxylin (Scale bar: 50 μm). Lung sections from 4 control patients and 5 SAVI patients were evaluated. Representative images are displayed on the left. For each stained slide, at least 5 images from the lung alveolar area were captured for quantification. Each dot in the quantification graph represents the average from different image measurements for an individual patient or control. The ratio of SMA/CD31 mean intensity was calculated using ImageJ. An increased SMA/CD31 ratio in the SAVI lung alveolar area suggests the role of EndMT. Data are represented as mean ±SEM. *$p < 0.05$ as determined by a two-tailed unpaired t-test.

(C) H&E staining in lung section from IPF and SAVI patients. The fibroblast foci is shown in IPF (left) but not shown in SAVI (right). Instead, thickened interlobular septa is observed in SAVI lung.

(D). Based on Masson's-Trichrome staining (left panel), different areas of mild (orange), moderate (red) and severe (green) fibrosis in Patient SAVI 8 lung tissue were selected on an adjacent slide stained for CODEX. At each condition, 6 individual areas were selected for CODEX quantification in Figure 1E.

(E) Immunofluorescent staining of endothelial and mesenchymal cell markers in lung alveola area from control and SAVI patients. Endothelial cell marker VE-Cadherin (VEcad, green) was co-stained with the mesenchymal marker smooth muscle actin (SMA, red) in lung tissue samples. Nuclei are stained blue with DAPI (Scale bar: 20 μm). White arrows indicate double positive cells in representative confocal images. The mean fluorescence pixel intensity for each channel was determined using the ZEN imaging software. The ratio of SMA/VEcad fluorescence pixel intensity was calculated from lung tissue staining of 5 control and 3 SAVI patients. Data are represented as mean ±SEM. *** $p < 0.001$ as determined by a two-tailed unpaired t-test.

#### Figure S2

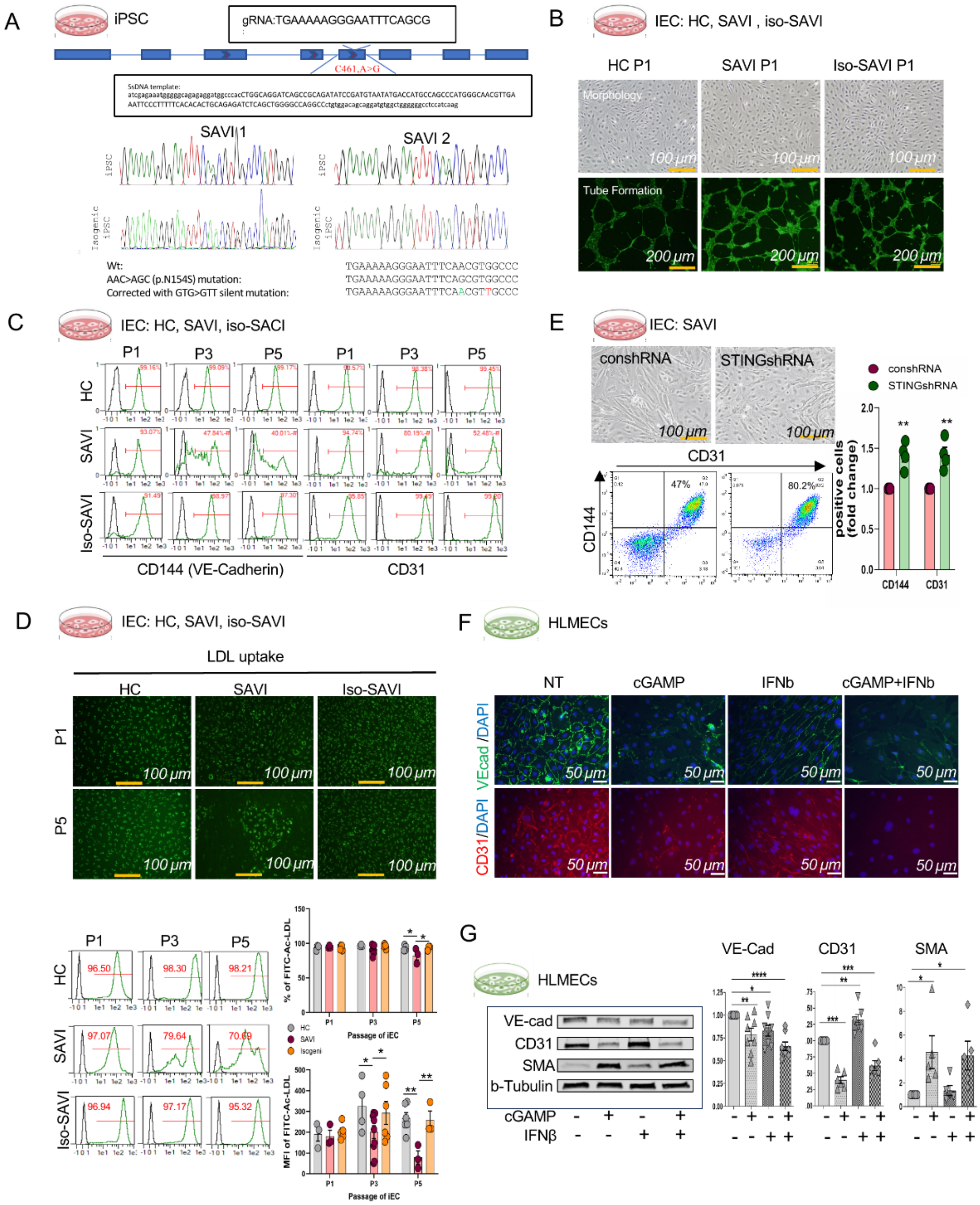


**Figure S2: iPSC-derived endothelial cells (iECs) from SAVI patients but not from isogenic and HC iECs spontaneously undergo EndMT upon cell passage. Related to Figure 2.**

(A) CRISPR/Cas9 correction of the *STING1* mutation (c.461A>G) in SAVI iPSCs .Guide RNA (gRNA) targeting STING1 and a donor oligo introducing a silent mutation were used to correct the heterozygous mutation (AAC to AGC). Sequencing confirmed correction and incorporation of the silent mutation (GTG to GTT). The sequence for gRNA is TGAAAAAGGGAATTTCAGCG. The STING1 mutation is highlighted in red. Exons are depicted as deep blue boxes with red arrows indicating the direction of transcription.

(B) Morphology and Tube Formation Assay in HC, SAVI, and iso-SAVI iECs at passage 1 (P1) displayed a similar cobblestone-like EC morphology and tube-forming capacity across all three groups at P1 (supplement to Fig 2B and 2D). Scale bars: 100 µm (top) and 200 µm (bottom).

(C) Flow cytometry profiles of endothelial surface marker CD144 (VE-cadherin) and CD31 at passage 1, 3 and 5 in HC, SAVI and iso-SAVI iECs. After CD31+ sorting, iECs were cultured to P1 and continued to be passaged until P5. At each passage, the presence of CD144 and CD31 positive cells was analyzed by flow cytometry. The quantification graph is shown in Fig 2C.

(D) Ac-LDL uptake at passages 1, 3, and 5. iECs were incubated with 10 µg/ml acetylated LDL (Ac-LDL) labeled with 1,1′-dioctadecyl-3,3,3′,3′-tetramethylindo-carbocyanine perchlorate (DiI-Ac-LDL). Fluorescence images were captured 4 hours after incubation at passages 1 and 5. Cells displaying green fluorescence had normal LDL uptake, which was observed in all groups except for SAVI iECs at passage 5. Flow cytometry profiles of iECs after incubation 24 hours from HC, SAVI, and iso-SAVI groups at passages 1, 3, and 5 are shown in the lower panel. The percentage of positive FITC-Ac-LDL cells and the Mean Fluorescence Intensity (MFI) in the FITC channel are summarized in the graph below. SAVI iECs demonstrated significantly lower FITC-Ac-LDL uptake function at P5, beginning at P3. Scale bar: 100 µm. Data are represented as mean ±SEM. **$p < 0.01$, *$p < 0.05$ as determined by 2-way ANOVA.

(E) STING knockdown rescues SAVI iEC phenotype. SAVI iECs were infected with control shRNA (conshRNA) or STING shRNA (STINGshRNA) at P1. By P5, the cell morphology had reverted from an elongated fibroblast-like shape back to a cobblestone EC-like shape, as shown in the upper panel. (Scale bar: 100 µm). EC surface markers CD144 and CD31 were analyzed by flow cytometry. Representative profiles indicate an increase in positive cells for CD144 and CD31 in the STINGshRNA group. The quantification graph is summarized from 3 individual SAVI patient iEC lines. Data are represented as mean ±SEM. **$p < 0.01$ determined by two-tailed unpaired t-test.

(F) Immunofluorescence staining of endothelial cell markers VE-Cadherin and CD31 in human lung microvascular endothelial cells (HLMECs) under STING activation. Cells were seeded into a 12-well plate at 60,000 cells per well. Cells were treated with or without 20 µg/ml cGAMP and/or 1000 units/ml IFNβ (human Interferon Beta 1a) for 5 days. VE-Cadherin (VE-Cad, green) and CD31 (red) were used as endothelial markers. Nuclei were stained with DAPI (blue). (Scale bar: 50 µm). Images are representative of n = 3 experiments. NT: no treatment.

(G) Western Blot analysis of EndMT in HLMECs. HLMECs were seeded into a 6-well plate at 100,000 cells per well. The cells were treated with or without 20 µg/ml cGAMP and/or 1000 units/ml IFNβ for 5 days. Cell proteins were collected and subjected to immunoblotting with endothelial markers (VE-cadherin (VE-Cad), CD31) and mesenchymal markers (alpha Smooth Muscle Actin (SMA)). Beta-tubulin served as a loading control. NT: no treatment. Representative images from 3 biological experiments are shown. Quantification graph normalized by loading control beta-tubulin. Data are represented as mean ± SEM. **** $p < 0.0001$, *** $p < 0.001$, ** $p < 0.01$, * $p < 0.05$ as determined by two-tailed unpaired t-test.

### Figure S3

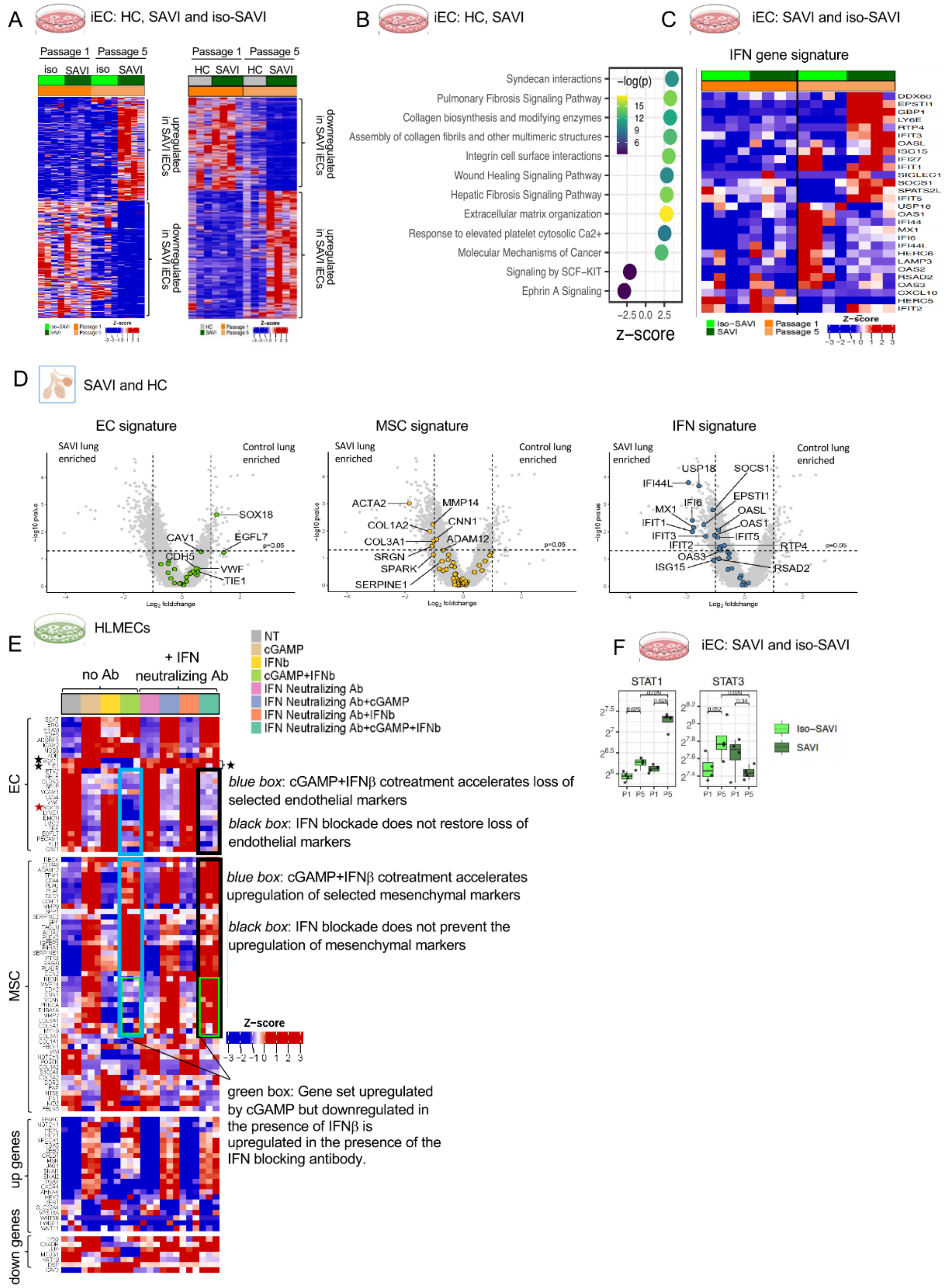

A
iEC: HC, SAVI and iso-SAVI
Passage 1
Passage 5
iso
SAVI
HC
upregulated in SAVI iECs
downregulated in SAVI iECs
B
iEC: HC, SAVI
Syndecan interactions
Pulmonary Fibrosis Signaling Pathway
Collagen biosynthesis and modifying enzymes
Assembly of collagen fibrils and other multimeric structures
Integrin cell surface interactions
Wound Healing Signaling Pathway
Hepatic Fibrosis Signaling Pathway
Extracellular matrix organization
Response to elevated platelet cytosolic Ca2+
Molecular Mechanisms of Cancer
Signaling by SCF−KIT
Ephrin A Signaling
−log(p)
z−score
C
iEC: SAVI and iso-SAVI
IFN gene signature
DDX60
EPSTI1
GBP1
LY6E
RTP4
IFIT3
OASL
ISG15
IFI27
IFIT1
SIGLEC1
SOCS1
SPATS2L
IFIT5
USP18
OAS1
IFI44
MX1
IFI6
IFI44L
HERC6
LAMP3
OAS2
RSAD2
OAS3
CXCL10
HERC5
IFIT2
Iso−SAVI
SAVI
Passage 1
Passage 5
Z−score
D
SAVI and HC
EC signature
MSC signature
IFN signature
SAVI lung enriched
Control lung enriched
SOX18
CAV1
EGFL7
CDH5
VWF
TIE1
ACTA2
MMP14
COL1A2
CNN1
COL3A1
ADAM12
SRGN
SPARK
SERPINE1
USP18
SOCS1
IFI44L
EPSTI1
MX1
IFI6
OASL
IFIT1
OAS1
IFIT3
IFIT5
IFIT2
RTP4
OAS3
ISG15
RSAD2
p=0.05
Log2 foldchange
-log10 pvalue
E
HLMECs
no Ab
+ IFN neutralizing Ab
NT
cGAMP
IFNb
cGAMP+IFNb
IFN Neutralizing Ab
IFN Neutralizing Ab+cGAMP
IFN Neutralizing Ab+IFNb
IFN Neutralizing Ab+cGAMP+IFNb
EC
MSC
up genes
down genes
blue box: cGAMP+IFNβ cotreatment accelerates loss of selected endothelial markers
black box: IFN blockade does not restore loss of endothelial markers
blue box: cGAMP+IFNβ cotreatment accelerates upregulation of selected mesenchymal markers
black box: IFN blockade does not prevent the upregulation of mesenchymal markers
green box: Gene set upregulated by cGAMP but downregulated in the presence of IFNβ is upregulated in the presence of the IFN blocking antibody.
Z−score
F
iEC: SAVI and iso-SAVI
STAT1
STAT3
Iso−SAVI
SAVI
P1
P5

**Figure S3: Multi-Modal profiling of SAVI iEC identifies transcriptional drivers of STING-mediated EndMT. Related to Figure 3.**

(A): Expression heatmap of differentially expressed genes between P5 and P1 iECs in the HC_SAVI and SAVi_iso-SAVI cohort from bulk RNAseq analysis.

(B) The most significant pathways, revealed by Ingenuity Pathway Analysis (IPA) of SAVI P5 vs SAVI P1 differentially expressed genes in the HC_SAVI cohort from bulk RNAseq analysis, are shown with positive z-scores indicating overall activation of pathway in SAVI P5. (supplementary to Fig3A).

(C) Expression heatmap of Type1 IFN gene signature between SAVI P5 and SAVI P1 iECs in the SAVI and SAVi_iso-SAVI cohort from bulk RNAseq analysis.

(D) The volcano plots of significantly changed genes in EndMT pathway gene signature and Type1 IFN gene signature in lung tissues from two control and one SAVI patient (SAVI 8) through GeoMx DSP Spatial Proteogenomic Assay.

(E) Bulk RNAseq analysis revealed expression heatmap of EndMT pathway genes in HLMECs under cGAMP or/and IFNβ treatment for 5 days with or without type 1 IFN neutralizing antibody (Ab).

(F) Bulk RNAseq analysis normalized gene expression of STAT1 and STAT3 in SAVI iEC_iso-SAVI iEC cohort at P1 and P5.

129 **Figure S4**

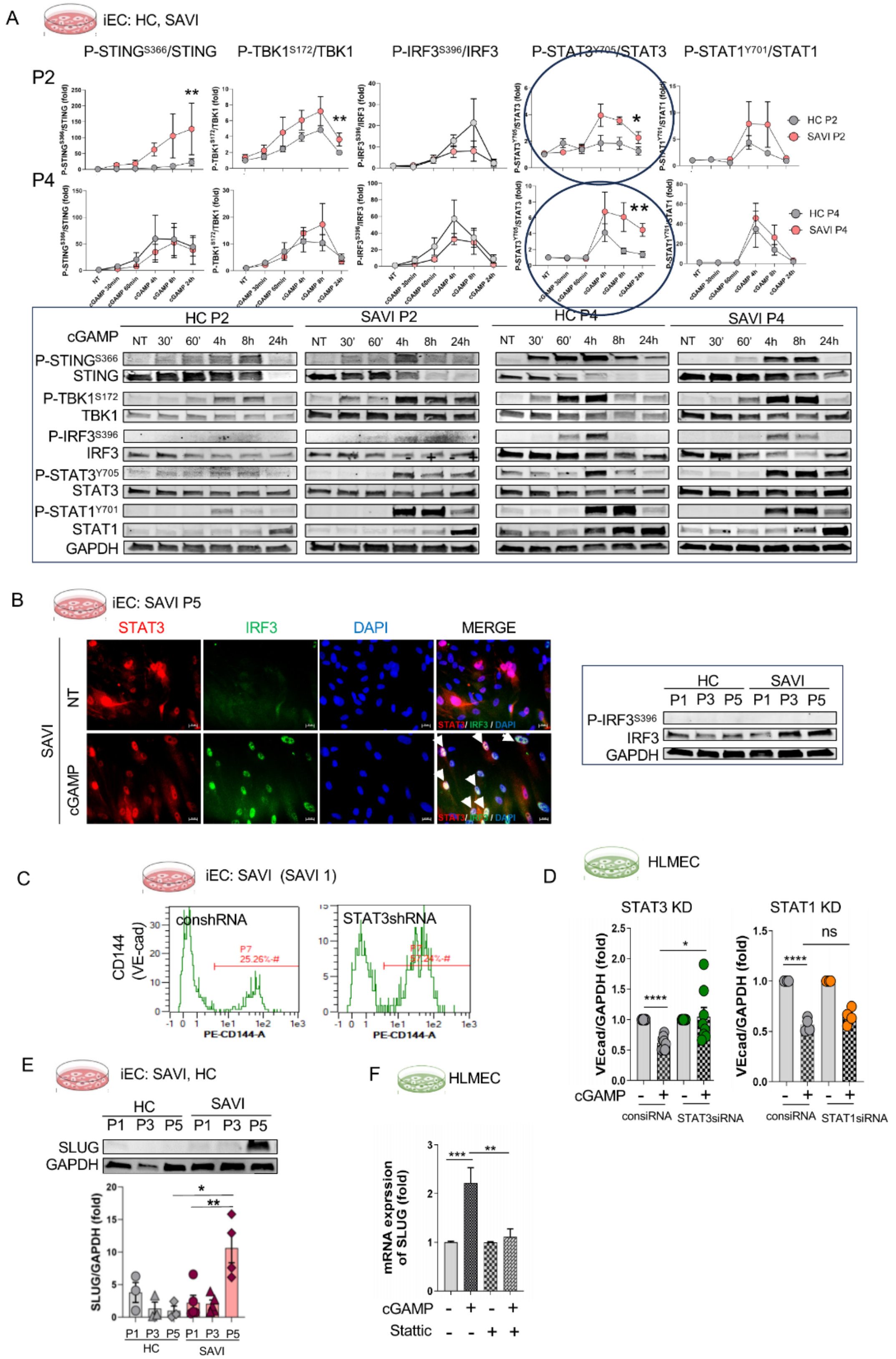


130

**Figure S4: Functional studies identify STAT3 as pivotal early driver of a SLUG dependent mesenchymal transcription program. Related to Figure 4.**

(A) cGAMP-Stimulated activation of the STING signaling pathway in HC and SAVI iECs**.** iECs from 3 individual HC and 3 individual SAVI patients were stimulated with 20 µg/ml 2'3'-cGAMP for 30 min, 60 min, 4 h, 8 h, and 24 h at P2 (early passage) and P4 (late passage). Protein expressions of P-STING$^{S366}$, P-TBK1$^{S172}$, P-IRF3$^{S396}$, P-STAT3$^{Y705}$, and P-STAT1$^{Y701}$ were normalized to total STING, TBK1, IRF3, STAT3, and STAT1 proteins. Time course curves are shown for P2 and P4 cells in both the HC and SAVI groups. Representative western blot images are shown below. Data are represented as mean ±SEM. **$p < 0.01$, *$p < 0.05$ as determined by 2-way ANOVA.

(B) IRF3-independent STAT3 activation in SAVI iECs. Left panel: Nuclear Translocation of STAT3 and IRF3 in SAVI iECs. Immunofluorescence staining of STAT3 (red) and IRF3 (green) was performed on iECs from one representative SAVI patient (SAVI1) at P5, with or without stimulation by 20 µg/ml cGAMP for 4 hours. White arrows indicate cells with co-localization of STAT3 and IRF3 in the nuclei, which were observed only in cGAMP-stimulated cells. Nuclei were stained with DAPI (blue). (Scale bar: 20 µm). Right panel: Western Blot images showing phosphorylation levels of P-IRF3$^{S396}$/IRF3 in HC iECs and SAVI iECs at passage 1, 3, and 5. GAPDH served as loading control.

(C) The rescue effect of STAT3 knockdown in SAVI iECs. SAVI iECs were infected with control shRNA (conshRNA) or STAT3 shRNA (STAT3shRNA) at P2. SAVI iECs at P5 were harvested for flow cytometry analysis. Representative profiles for CD144 and CD31 in SAVI1 patient iEC line. shown here serve as a supplement to Fig4E.

(D) The effect of knockdown of STAT3 and STAT1 on cGAMP/IFNβ-Induced downregulation of endothelial cell markers VE-Cadherin in HLMECs. HLMECs cells were transfected with silencing RNA (siRNA) targeting control, STAT3, or STAT1 for 48 hours, followed by treatment with 20 µg/ml 2'3'-cGAMP and 1000 units/ml IFNβ for 3 days. Protein samples were then harvested for Western blot analysis of the endothelial markers VE-Cadherin (VEcad) with GAPDH serving as the loading control. The quantification graph is summarized from 4-8 biological experiments. Data are represented as mean ±SEM. **** $p < 0.0001$, *** $p < 0.001$, **$p < 0.01$, *$p < 0.05$ as determined by a two-tailed unpaired t-test.

(E) Dynamic increased SLUG protein expression in SAVI iECs. Protein samples were collected from iECs derived from 3 individual HC and 3 individual SAVI patients at passages 1, 3, and 5. Western blot analysis for SLUG was conducted. GAPDH was used as a loading control. Quantification graphs and representative western blot images are shown. Data are represented as mean ±SEM. **$p < 0.01$, *$p < 0.05$ as determined by a two-tailed unpaired t-test. Related to Figure 4F.

(F) The effect of STAT3 inhibitor on 2'3'-cGAMP-induced mRNA expression of *SNAI2*/SLUG in HLMECs. HLMECs were treated with or without 20 µg/ml cGAMP for 24 hours. The STAT3 inhibitor, Stattic (5 µM), was pre-treated for one hour. DMSO served as a vehicle control. RNA was extracted for RT-PCR at 24 hours (day 1). n = 4. Data are represented as mean ±SEM. *** $p < 0.001$, **$p < 0.01$ as determined by a two-tailed unpaired t-test.

### Figure S5

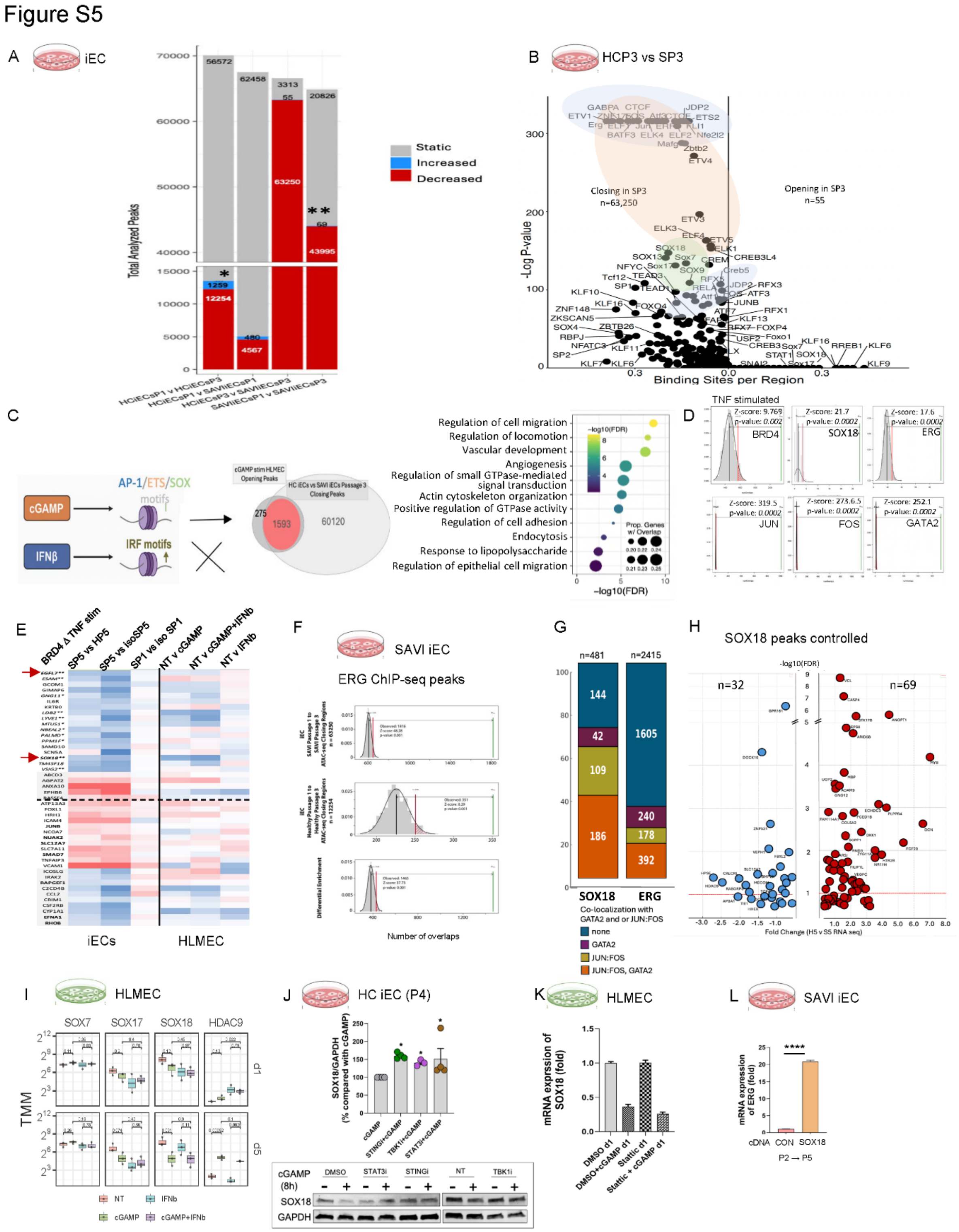
Figure S5
A
iEC
Total Analyzed Peaks
Static
Increased
Decreased
HCiECsP1 v HCiECsP3
HCiECsP1 v SAVIiECsP1
HCiECsP3 v SAVIiECsP3
SAVIiECsP1 v SAVIiECsP3
B
HCP3 vs SP3
Closing in SP3
n=63,250
Opening in SP3
n=55
-Log P-value
Binding Sites per Region
C
cGAMP
IFNβ
AP-1/ETS/SOX
motifs
IRF motifs
Regulation of cell migration
Regulation of locomotion
Vascular development
Angiogenesis
Regulation of small GTPase-mediated signal transduction
Actin cytoskeleton organization
Positive regulation of GTPase activity
Regulation of cell adhesion
Endocytosis
Response to lipopolysaccharide
Regulation of epithelial cell migration
-log10(FDR)
D
TNF stimulated
BRD4
SOX18
ERG
JUN
FOS
GATA2
E
BRD4 Δ TNF stim
SP5 vs HP5
SP5 vs isoSP5
SP1 vs iso SP1
NT v cGAMP
NT v cGAMP+IFNb
NT v IFNb
iECs
HLMEC
F
SAVI iEC
ERG ChIP-seq peaks
Number of overlaps
G
n=481
n=2415
SOX18
ERG
Co-localization with GATA2 and or JUN:FOS
none
GATA2
JUN:FOS
JUN:FOS, GATA2
H
SOX18 peaks controlled
n=32
n=69
Fold Change (H5 v S5 RNA seq)
I
HLMEC
SOX7
SOX17
SOX18
HDAC9
TMM
NT
IFNb
cGAMP
cGAMP+IFNb
J
HC iEC (P4)
SOX18/GAPDH (% compared with cGAMP)
cGAMP (8h)
DMSO
STAT3i
STINGi
NT
TBK1i
SOX18
GAPDH
K
HLMEC
mRNA exprssion of SOX18 (fold)
DMSO d1
DMSO+cGAMP d1
Static d1
Static + cGAMP d1
L
SAVI iEC
mRNA expression of ERG (fold)
cDNA
CON
SOX18
P2 → P5

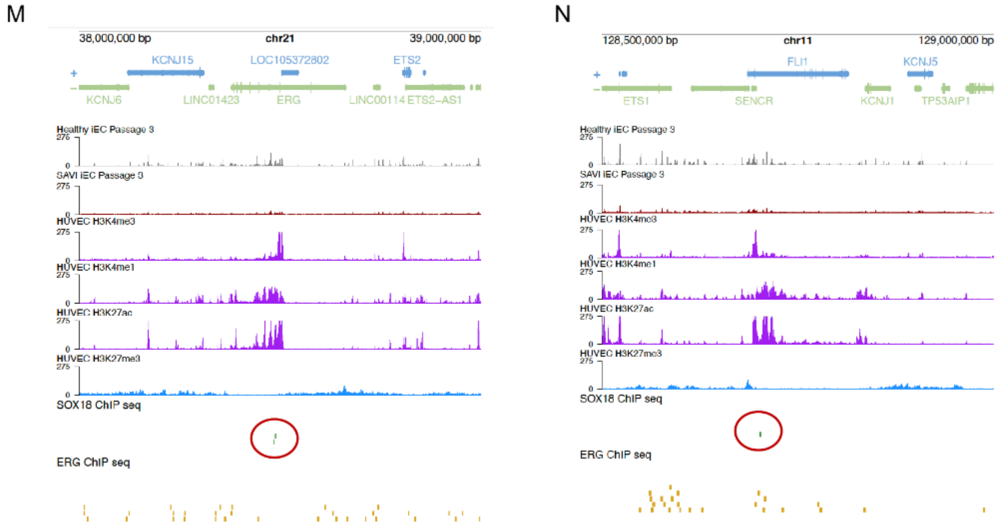


**Figure S5: STING-driven enhancer remodeling targets GATA2/AP-1/SOX18 networks that maintain endothelial stability. Related to Figure 5.**

(A) Histogram of static and dynamically changing chromatin regions in the iECs. Blue are regions becoming accessible and red are region that are closing. SAVI iECs (*) had a 3-fold change in accessible chromatin regions (CRs) with mostly reduced accessibility compared to the difference in healthy control (**) P1 and P3 iECs (44,064 vs. 13,513 CRs; *$p<0.0001$, by Fisher exact).*

(B) Motif enrichment (FIMO) in regions becoming accessible in HC P1 vs HC P3 in iECs: This volcano plot shows the enrichment of transcription factor binding motifs within the differentially accessible regions between HC iEC ATAC-seq libraries from P1 and P3 (left volcano plot) and HC and SAVI patient iECs are P3 (SP3) (right volcano plot). Enrichment of binding motifs with increased accessibility are on the right and with decreased accessibility on the left. Blue clouds denote mostly AP-1 TF family binding motives, rose clouds denote ETS-motif containing TF motif binding motifs and green clouds denote SOX family member TF binding motifs.

(C) Schematic showing that cGAMP stimulation in HLMECs increases accessibility of AP-1, ETS motif containing-, and SOX motif containing TFs in contrast IFNβ stimulation expose CRs with predominantly IRF motives. The Euler plot shows cGAMP-responsive regions (1,593 of 1,868 total chromatin regions from Figure 5D-E) that overlap with regions selectively losing accessibility in SAVI iECs transitioning to a mesenchymal phenotype). Pathway analysis of genes within these regions (right) revealed enrichment for processes involved in endothelial integrity and vascular homeostasis in the cGAMP stimulated samples in contrast to the IFNβ stimulated HLMECs.

(D) ChIP-seq motifs from HUVEC data were mapped to the 1593 STING-responsive overlapping regions and showed significant enrichment in inflammatory BRD4, SOX18, GATA2 and JUN and FOS ChIP-seq marks.

(E) Inflammatory BRD4 redistribution regulated genes: Pattern of transcriptional regulation in SAVI iECs undergoing EndMT and in HLMECs stimulated with cGAMP follows that in TNF-induced BRD4 redistribution mediated transcriptional changes shows downregulation of endothelial identity genes in regions losing BRD4 marks, including *EGFL7, ESAM, LYVE1,*

*SOX18, LDB2, VSIG2 (indicated with* **), and genes regulating endothelial stability or antiproliferative and cytoskeleton regulation including *GNG11,MTUS1, NBEAL2, PALMD, PPM1F, TM4SF18 (indicated by *)* in cGAMP stimulated HLMECs at day 5 and SAVI iECs passage 5 vs HC iECs passage 5, SAVI iECs vs iso-SAVI iECs, but not in SAVI iECs passage 1 compared to HC iECs passage 1. Upregulated genes are in regions with gained BRD4 chip seq marks. Red arrows point to *SOX18* and *EGFL7* which are also downregulated in SAVI lung tissue (Figure S3D).

(F) These plots show the results of permutation tests from the regioneR package for the overlap between ERG ChIP-seq peaks from Kalna et al. HUVEC dataset and the SAVI P1 to SAVI P3 closing regions, or the HC P1 to HC P3 closing regions. A differential permutation test was also conducted to compare the relative enrichment of ERG ChIP-seq peaks in these two sets of regions. The green bar indicates the observed number of overlaps between the datasets. The black bar indicates the mean value of overlaps between the transcription factor ChIP-seq peaks and the random permuted regions. the distribution of overlaps from the permutations are shown as the gray histogram). The red bar indicates the number of overlaps at the threshold of significance $p = 0.05$. Differential enrichment analysis (bottom plot) showed significant overrepresentation of ERG binding sites in SAVI closing CRs at P3 (1465 regions, $z = 37.73$ ; $p < 0.001$).

(G) SOX18 preferentially colocalizes with AP-1/GATA2 enhancer elements compared with ERG: SOX18 ChIP-seq peaks showed significantly greater co-localization with AP-1 and/or GATA2 compared to ERG peaks (two-tailed $\chi^2$ with Yates' correction, $p < 0.0001$), consistent with SOX18 marking enhancer domains more prominently than ERG.

(H) Volcano plot showing differential gene expression in SAVI iECs v. HC iEC at passage 5, associated with SOX18 CHIP-seq peaks (n= 113 regulating 101 genes) in all CR regions that become inaccessible in SAVI iECs passage 3. 42 SOX18 CHIP-seq peaks were associated with 32 downregulated genes (blue circles) (RNA seq at performed at passage 5) and 71 were associated with 69 upregulated genes (red circles).

(I) RNAseq analysis of normalized gene expression of SOX7, SOX17, SOX18 and HDAC9 in nontreated, cGAMP or IFNβ or both cGAMP+IFNβ stimulated HLMECs after 1 and 5 days of stimulation.

(J) The effect of STING/TBK1/STAT3 inhibitors on 2'3'-cGAMP-induced SOX18 downregulation in HC iECs. iECs derived from 3 individual HC patients were treated with 20 μg/ml 2'3'-cGAMP for 8h at passage 4. STING, TBK1 and STAT3 inhibitors were pre-treated for one hour ahead. Compared to 2'3'-cGAMP alone, both STING inhibitor IFM35883 (STINGi, 2.5 μM), TBK1 inhibitor MRT67307 (TBK1i, 5 μM) and STAT3 inhibitor (STAT3i, 5 μM) significantly rescued (increased by about 50%) cGAMP-induced SOX18 downregulation. Quantification graphs and representative western blot images are shown. Data are represented as mean ±SEM. n = 4. *$p < 0.05$ as determined by Mann-Whitney test.

(K) The effect of STAT3 inhibitor (Stattic) on 2'3'-cGAMP-induced mRNA downregulation of SOX18 in HLMECs. HLMECs were treated with or without 20 μg/ml cGAMP for 24 hours. The STAT3 inhibitor, Stattic (5 μM), was pre-treated for one hour. DMSO served as a vehicle control. RNA was extracted for RT-PCR at 24 hours (day 1). Data are represented as mean ±SEM (two biological experiments, three technical replicates each).

(L) SOX18 overexpression upregulated mRNA expression of ERG in SAVI iECs at P5. SAVI iECs were transduced at P2 with control or SOX18 cDNA; RNA was extracted for RT-PCR at P5. Data are represented as mean ± SEM; ****$p < 0.0001$, two-tailed unpaired t-test. Experiments were conducted in SAVI1 iEC line for 2 individual biologicals with 6 technical repeats.

(M) and (N): Genome tracks show SOX18 binding at intragenic enhancer elements (red circle) marked by H3K4me1 and H3K27ac within the ERG (M) and FLI1 (N) loci, consistent with active endothelial lineage-maintenance enhancers.

**Figure S6**

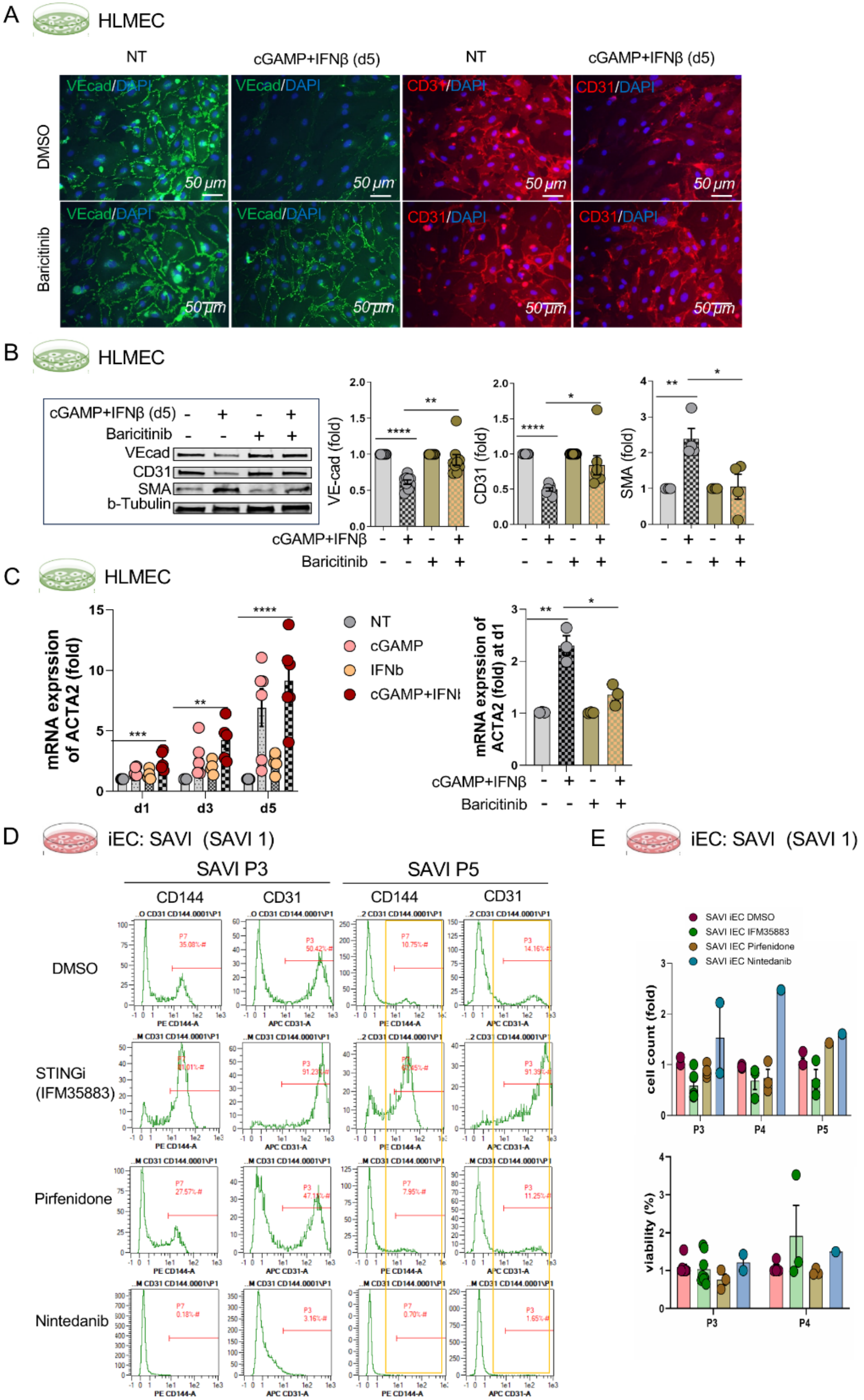
A HLMEC
NT
cGAMP+IFNβ (d5)
DMSO
Baricitinib
VEcad/DAPI
CD31/DAPI
50 μm
B HLMEC
cGAMP+IFNβ (d5)
Baricitinib
VEcad
CD31
SMA
b-Tubulin
VE-cad (fold)
CD31 (fold)
SMA (fold)
cGAMP+IFNβ
Baricitinib
C HLMEC
mRNA exprssion of ACTA2 (fold)
d1
d3
d5
NT
cGAMP
IFNb
mRNA exprssion of ACTA2 (fold) at d1
D iEC: SAVI (SAVI 1)
SAVI P3
SAVI P5
CD144
CD31
DMSO
STINGi (IFM35883)
Pirfenidone
Nintedanib
PE CD144-A
APC CD31-A
E iEC: SAVI (SAVI 1)
SAVI iEC DMSO
SAVI IEC IFM35883
SAVI IEC Pirfenidone
SAVI iEC Nintedanib
cell count (fold)
viability (%)
P3
P4
P5

**Figure S6: STING inhibition but not drugs approved for treatment of pulmonary fibrosis preserve the endothelial phenotype in STING induced EndMT. Related to Figure 6.**

(A) The effects of the JAK/STAT inhibitor Baricitinib on immunofluorescence staining of endothelial cell markers VE-Cadherin and CD31 in HLMECs under STING activation. HLMECs cells were treated with 20 μg/ml 2'3'-cGAMP and 1000 units/ml IFNβ along with either Baricitinib (1 μM) or DMSO control for 5 days. Immunofluorescence staining displayed endothelial markers VE-Cadherin (VE-Cad, green) and CD31 (red), with nuclei stained with DAPI (blue) (Scale bar: 50 μm). Treatment with cGAMP/IFNβ led to the loss of endothelial markers. Baricitinib partially restored EC cell-cell junctions, particularly VE-Cadherin. Images are representative of n = 3 experiments. NT: no treatment.

(B) The effects of the JAK/STAT inhibitor Baricitinib on cGAMP/IFNβ-Induced EndMT in HLMECs. HLMECs cells were treated with 20 μg/ml cGAMP and 1000 units/ml IFNβ along with either Baricitinib (1 μM) or DMSO control for 5 days. Protein samples were harvested on day 5 for Western blot analysis. The relative abundance of endothelial markers VE-Cadherin (VEcad) and CD31, as well as the mesenchymal marker SMA in each group, is displayed on the right. β-Tubulin served as the loading control. Baricitinib significantly reduced cGAMP/IFNβ-induced EndMT. Representative images are shown on the left. The quantification graph is summarized from 3 biological experiments. Data are represented as mean ±SEM. **** $p < 0.0001$, **$p < 0.01$, *$p < 0.05$ as determined by two-tailed unpaired t-test.

(C) The effect of JAK/STAT inhibitor baricitinib on cGAMP/IFNβ-induced mRNA expression of ACTA2 (SMA) in HLMECs at an day 1. HLMECs cells were treated with or without 20 μg/ml cGAMP and/or 1000 units/ml IFNβ for 1, 3, and 5 days. RNA was extracted for RT-PCR. mRNA expression of ACTA2 (SMA), normalized by the internal control 18S, was measured at different time points. Starting from day 1, cGAMP/IFNβ significantly increased ACTA2 (SMA) mRNA expression. Baricitinib (1 μM) was pre-treated for one hour. DMSO served as a vehicle control. RNA was extracted for RT-PCR at 24 hours (day 1). cGAMP/IFNβ-induced upregulation of mRNA expression of ACTA2 (SMA) was significantly restrained by Baricitinib. Data are summarized from 3 biological experiments. Data are represented as mean ±SEM. **** $p < 0.0001$, *** $p < 0.001$, **$p < 0.01$ as determined by a two-tailed unpaired t-test.

(D) Representative flow cytometry profiles of treatments with DMSO control, IFM35883 (STINGi), Pirfenidone, and Nintedanib in SAVI iECs. The treatments of STING inhibitor IFM35883 (2.5 μM), Pirfenidone (10 μM), Nintedanib (1 μM), and vehicle control DMSO were initiated at passage 2 in SAVI iECs. At each passage, CD144-positive cells and CD31-positive cells were analyzed by flow cytometry. Representative profiles for CD144 and CD31 at passages 3 and 5 are shown here as a supplement to Fig6C.

(E) The effects of STING Inhibitor IFM35883, Pirfenidone, and Nintedanib on cell number and viability in SAVI iECs. The treatments of STING inhibitor IFM35883 (2.5 μM), Pirfenidone (10 μM), Nintedanib (1 μM), and vehicle control DMSO were initiated at passage 2 in SAVI iECs. At each passage, cell number and viability were measured using an automatic cell counter with Trypan blue. Data are represented as mean ± SEM.

Table S1.

| SAVI # (n=13) | STING1 variant | Histo H&E (n=7) | Histo CD31/SMA (n=5) | IF CD31 (n=3) | IF VEcad/SMA (n=3) | IF pSTAT3 (n=3) | IECs (n=4) | isoIECs (n=2) | GeoMx (n=1) | CODEX (n=1) | Fibroblast cell line (n=6) | Chest CT (n=13) |
|---|---|---|---|---|---|---|---|---|---|---|---|---|
| **Radiographic Lung Disease** | | | | | | | | | | | | |
| **SAVI 1** | N154S | X | X | X | X | X | X | X | nd | nd | nd | X |
| **SAVI 2** | N154S | nd* | na | na | na | na | X | X | na | na | nd | X |
| **SAVI 5** | N154S | X | X | X | X | X | na | na | nd | nd | nd | X |
| **SAVI 6** | N154S | X | nd | nd | nd | nd | na | na | nd | nd | nd | X |
| **SAVI 7** | V147L | X | X | nd | nd | nd | na | na | nd | nd | X | X |
| **SAVI 8** | V155M | X | X | X | X | X | na | na | X | X | nd | X |
| **SAVI 9** | V155M | X | X | nd | nd | nd | na | na | nd | nd | nd | X |
| **SAVI 10** | V155M | X | nd | nd | nd | nd | na | na | nd | nd | X | X |
| **SAVI 11** | N154S | nd* | na | na | na | na | na | na | na | na | X | X |
| **SAVI 13** | R281W_hm | nd* | na | na | na | na | na | na | na | na | X | X |
| **No Radiographic Evidence of Lung Disease** | | | | | | | | | | | | |
| **SAVI 3** | N154S | nd* | na | na | na | na | X | nd | na | na | nd | X |
| **SAVI 4** | V147L | nd* | na | na | na | na | X | nd | na | na | X | X |
| **SAVI 12** | H72N | nd* | na | na | na | na | na | na | na | na | X | X |

Table S1: SAVI Patient Specimen Information: Hm, homozygous variant; histo, histopathology; IF, immunofluorescence; SMA, smooth muscle actin; VEcad, vascular endothelial cadherin; IECs, induced pluripotent stem cells (iPSC) derived endothelial cells; IsoiECs, isogenic iPSC derived endothelial cells; CT, computed tomography; X, performed; nd, not done; na: material not available; *, These patients did not have a lung biopsy performed.
